# Movement execution defines a distinct neural state in dyskinesia and enhances decoding

**DOI:** 10.1101/2025.06.09.658641

**Authors:** J. Habets, T. Merk, V. Mathiopoulou, J. Kaplan, R. Lofredi, J. L. Busch, T. Binns, R. Köhler, A. Memarpouri, K. Faust, P. Krause, GH. Schneider, WJ. Neumann, P. Tovote, A. Kühn

## Abstract

Parkinson patients suffer from levodopa-induced dyskinesia, which occur adversely to chronic dopaminergic treatment. These abnormal involuntary movements can only partly be actively suppressed and affect quality of life. A lowered motor inhibition during hyperdopaminergic states, associated with structural and plasticity changes in the cortico-basal-ganglia motor network, is hypothesized to enable dyskinesia. Oscillatory cortico-subthalamic patterns associated with dyskinesia are proposed as adaptive neuromodulation biomarkers, but their dependence on behavioral states such as dyskinetic movement presence or suppression remains unknown.

We studied cortico-subthalamic oscillations in 22 Parkinson’s patients during dyskinesia-evoking protocols. We clinically differentiated between non-dyskinetic and dyskinetic periods, and defined movement presence with kinematics, leading to four behavioral states containing rest, voluntary movements, movement suppression during dyskinesia, and dyskinetic movements.

Elevated subthalamic theta-activity and attenuated beta-activity was found during both dyskinetic movement suppression and execution, while cortico-subthalamic gamma-activity only increased during dyskinetic movement execution. Subthalamic spectral changes significantly predicted dyskinesia presence, and movement presence significantly affected the predictive performance. A movement-aware classifier enhanced dyskinesia detection based on movement-depending biomarkers containing cortical oscillations and gamma-bands.

We propose movement execution during dyskinesia to be a distinct behavioral and neural microstate within a dopamine-depending dyskinetic macrostate, that can enhance dyskinesia classification for adaptive neuromodulation.

## Main

Dopaminergic replacement therapy is the cornerstone to treat hypokinetic symptoms in Parkinson’s disease (PD), but also adversely causes disabling abnormal involuntary movements, defined as levodopa induced dyskinesia (LID) ^1–4^. As disease progression continues and requires larger and more frequent dopaminergic intakes, LID occurs in up to 80% of PD patients, impacting quality of life ^1,3,5^. LID primarily emerges during moments of dopaminergic peaks and has a fluctuating heterogeneous clinical phenotype of involuntary movements, alternating with episodes of voluntary motor suppression and subjective internal motor restlessness ^2^. Typically, LID is treated with either pharmacological interventions that improve levodopa’s efficacy and allow dopamine reduction ^4^ or target specific striatal projecting neurotransmitter-receptors ^6^; or with device-aided therapies such as deep brain stimulation (DBS) ^5,7–10^. Although overstimulation with DBS can induce dyskinesia ^11^, DBS generally alleviates LID by allowing a reduction of dopaminergic medication ^12^ and possibly by a long-term desensitization of dopamine receptors ^13^.

Current understandings of the pathophysiology of LID hypothesize that a reduced cortical long-term depotentiation alters cortico-striatal synaptic plasticity ^14,15^. The impaired synaptic plasticity fails to balance dopaminergic cortico-basal-ganglia excitation and increases the likelihood of movement ^16^. On a striatal level, chronic dopamine depletion is hypothesized to cause compensatory structural changes in striatal projection neuron architecture and fast spiking interneurons ^17–19^. These compensatory neural changes shift the balance between the canonical direct and indirect motor pathways to restore homeostasis during hypo-dopaminergic periods ^17,18,20,21^. Moreover, long term dopaminergic depletion leads to abnormal spiking patterns of SPN neurons with loss of spatial and temporal selectivity ^22^. Neural dynamics can partially be restored by dopaminergic treatment, but during hyper-dopaminergic periods, inverse abnormalities in striatal activity pattern with overcompensated reduction in motor inhibition may occur, facilitating LID ^22–26^. Since the subthalamic nucleus (STN) is specifically involved in movement planning, execution, and cancellation ^27–29^, STN DBS can directly modulate physiological motor stopping ^30^, but also dyskinetic movement ^11^.

Invasive neurophysiological studies explored biomarkers that represent spectral network oscillations during different motor, cognitive, and emotional states in PD ^31^. Subthalamic beta activity has been established as a symptom-specific biomarker for bradykinesia ^28^, independent of disease ^32,33^. Furthermore, cortico-subthalamic gamma activity has been used to decode movement initiation, velocity and stopping ^34–36^, whereas increased theta band activity has been described in conflicting cognitive tasks during decision making and in impulse control disorders ^37,38^. Regarding LID, elevated subthalamic and cortical gamma-band (60 – 90 Hz) activity has been associated with LID presence and severity ^39–41^, similar to elevated subthalamic theta- and alpha-band (resp. 4 – 8 and 8 – 12 Hz) activity ^37^, and decreased subthalamic beta-band (13 – 35 Hz) activity ^40,42^. The spatiotemporal patterns of oscillations associated with LID so far, have not been differentiated with respect to the highly variable behavioral phenotype of LID ^2^, i.e. movement execution and movement suppression, and therefore their behavioral relevance within LID remains unclear ^31^. This highly variable behavioral pattern within LID, containing both presence and absence of movement execution ^2^, may modulate neural oscillations by the degree of successful active motor suppression ^43^. Moreover, recent rodent studies could show that distinct dyskinetic movements are associated with distinct neural patterns ^21^. In this vein, we hypothesize that distinct behavioral states of movement absence and presence within dyskinetic periods are associated with distinct neural oscillatory pattern that can be used as neurophysiological biomarkers.

For this concept, we applied macro- and microstates that were recently introduced to explain integrated systemic, cardio-behavioral states in the context of emotional challenge ^44^. They allow the characterization of temporal dynamics within behavioral and physiological changes over variable time-periods ^44^. Herein, slow transitions are termed as macrostate changes and contrast fast-changing, momentary microstates, which reflect short-lasting distinct behavioral or neural patterns. Our hypothesis includes a slow, macrostate transition from the hypodopaminergic OFF state to the hyperdopaminergic ON state in PD, which lowers motor inhibition based on the compensatory basal ganglia changes due to chronic dopamine depletion. On the background of the hyperdopaminergic ON state, fast-changing microstates may occur that lead to the execution of dyskinetic movements. To investigate the hypothesized states-concept, we studied invasive subthalamic and cortical neural oscillations from the hypodopaminergic OFF state to the hyperdopaminergic ON state with LID, during the execution of voluntary and dyskinetic movements and during dyskinetic active movement suppression. We identified distinct subthalamic theta and beta oscillations associated with the slow-changing dopaminergic periods considered as a macrostate. Fast-changing patterns of cortico-subthalamic gamma oscillations that depend on the presence of dyskinetic movement could be defined as microstate changes. The movement-depending character of these spectral biomarkers was utilized to develop a movement-aware LID detection model. Our results demonstrate that including this behavioral context of movement presence enhances the decoding of LID using models based on only-cortical oscillations and using models based on only-gamma-band biomarkers. The identification of the behavioral relevance of these LID-associated oscillations can help us to better understand LID’s pathophysiology and motor control in general, and to translate cortico-subthalamic biomarkers into robust clinical adaptive DBS strategies.

### Dyskinetic periods contain moments with and without objectified movement execution

To describe LID-related spectral patterns, we included twenty-one PD patients implanted with bilateral STN DBS electrodes, of which thirteen received a temporary unilateral electrocorticography (ECoG) electrode (mean age: 61 years, standard deviation (sd): 9.2 years, mean disease duration: 10 years (sd: 4 years)) (see Table S1). One additional subject was included to allow an external validation analysis and was not included in descriptive analyses and not in cross-validation model development (see Table S1). All participants performed a dyskinesia-provoking experiment during the interval (two to seven days) between their surgical implantation of DBS-electrodes and its connection to the internal pulse generator ^45^. Participants started the experimental protocol in the medication-OFF condition (at least 12 hours of dopamine withdrawal) followed by a supra-threshold dosage of fast-acting levodopa (circa 1.5 morning dose of Madopar LT ®, Roche Pharma). STN and cortical activity were recorded during repetitive rest and unilateral self-paced hand tapping tasks, intermingled with unscripted pauses to avoid fatigue and experiment abortion (see Figure 1 and Fig. S1).

**Figure 1:**
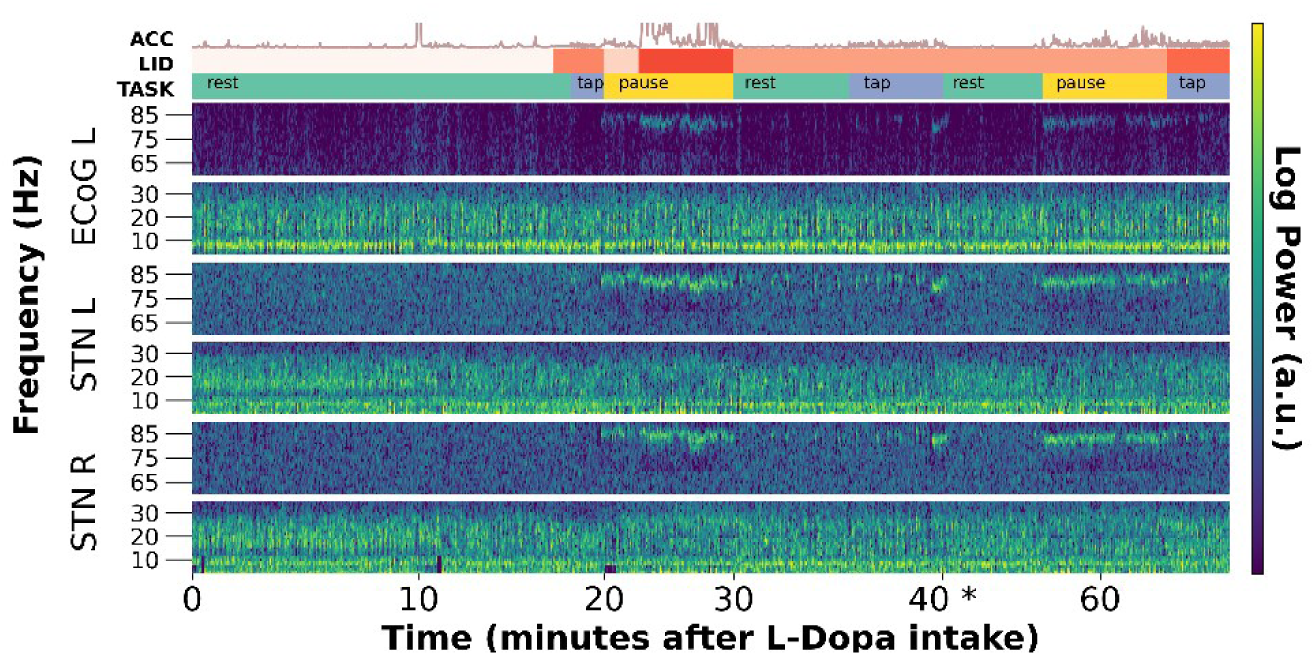
Single subject example of spectral changes and behavioral heterogeneity during dyskinetic periods. Neural, behavioral, and clinical data from one subject is shown. The first upper row (ACC) represents the participant’s activity, recorded via bilateral accelerometers placed on the index-fingers. The line shows the average of the z-scored root mean squares of both accelerometers. The second upper row (LID) represents the clinically assessed dyskinesia presence and severity (defined as sum CDRS) over time. The first white part represents the non-dyskinetic period. A dyskinetic period is represented by different shades of orange, where darker orange represents more severe LID. LID occurred bilateral in this subject, with more severe dyskinetic movements in the right body side compared to the left. The third upper row represents the task performed (TASK) during the experiment, in which *rest* contained seated rest*, tap* contained a seated self-paced, unilateral hand-tapping task, and *pause* contained an unscripted pause for the participant to prevent fatigue. The three neural time-frequency plots represent spectral unilateral cortical (ECoG) and bilateral subthalamic (STN) activity. Note the decrease in beta activity bilaterally in the STN at about 10 minutes after levodopa intake and during LID, and irregular gamma synchronization during LID, mostly in the presence of movement. *: The recording was interrupted around minute 45 for a toilet break and the x-axis is adjusted accordingly. ECoG: electrocorticography, Hz: Hertz, L-Dopa: levodopa, LID: levodopa induced dyskinesia, STN: subthalamic nucleus.

A clinical movement disorders specialist assessed the presence and severity of LID based on video recordings, using the Clinical Dyskinesia Rating Scales (CDRS). ^46^ The presence of LID was observed in eighteen out of twenty-one participants (86%), of which fifteen had a preoperative history of LID (83%) (see Methods). One participant, that did not develop LID, had a preoperative LID history. The eighteen patients that did develop LID had mean peak dyskinesia severities of 5 CDRS points (sd: 3, range: 2 to 13 CDRS points), out of a maximum of 28 points (the latter corresponds to severe dyskinetic movement in all seven rated body parts).

The distributed repetitions of resting and tapping tasks blocks led to the recording of movement presence and movement absence in both dyskinetic states and non-dyskinetic states (Figure 1). We defined a dyskinetic period as a time interval that starts at the first observed dyskinetic movement and prolongs as long as the patient regularly shows abnormal involuntary movements. Dyskinetic periods will also contain moments of suppression of involuntary movements, usually achieved by active movement suppression with voluntary muscular co-contraction in between dyskinetic movements (Figure 2). Fluctuations in the clinically assessed severity of a dyskinetic period were assessed on a minute level, while the observation of actual movement was objectively assessed on a sub second level with accelerometer data. Consequently, we differentiated between 1) movement presence during the non-dyskinetic period (i.e. voluntary movements), 2) movement absence during the non-dyskinetic period (i.e. the resting state), 3) movement presence during the dyskinetic period (i.e., execution of both voluntary and involuntary movements, during episodes clinically confirmed as dyskinetic), and 4) movement absence during the dyskinetic period (i.e., epochs of motor suppression, lacking objectified movement execution, within periods clinically confirmed as dyskinetic). This behavioral concept, that dyskinetic periods contain epochs of both movement presence and absence, is essential to assess different neural microstates, i.e. dyskinetic movements, during the hyperdopaminergic macrostate characterized by a dyskinetic period.

**Figure 2:**
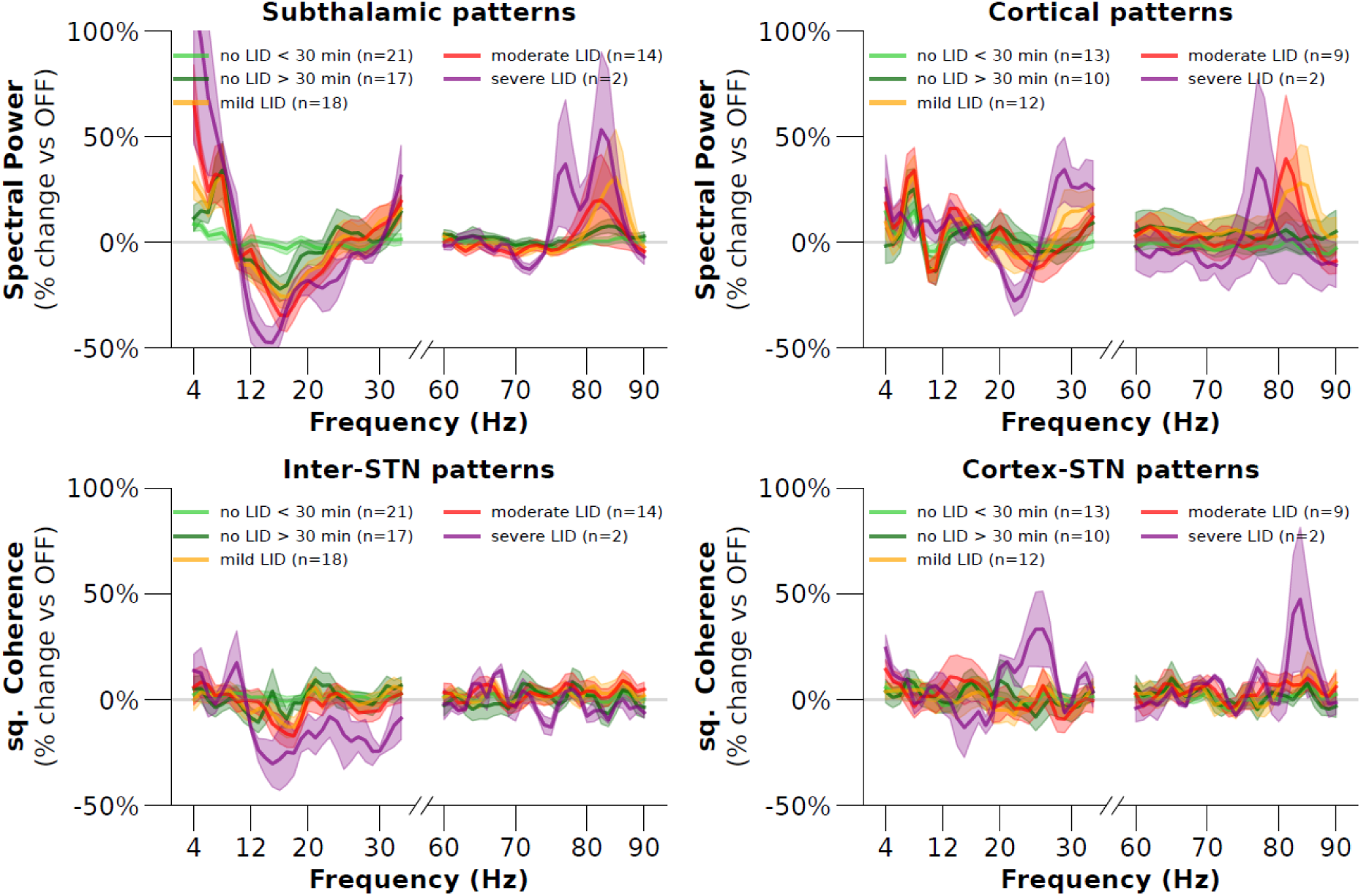
Spectral subthalamic and cortical patterns change during dyskinetic periods depending on dyskinesia severity. All four panels show the changes in spectral patterns as percentage changes relative to the dopaminergic OFF condition at rest up to five minutes after levodopa-intake. Three grades of dyskinesia severity are visualized: mild LID in yellow (1 to 3 CDRS points), moderate in red (4 to 7 CDRS points), and severe in purple (8 or more CDRS points). The OFF and ON conditions are visualized in light and dark green. The numbers in the legend represent the number of subjects that contributed neural data to a category. The upper row shows power spectral densities of subthalamic nucleus (STN) (upper left) and cortical (upper right) recordings. Note the increase in STN theta and decrease in beta activity that scale with severity of dyskinesia, the scaling effect is less prominent for gamma synchronization with dyskinesia in STN and cortex. Spectral powers are calculated using Welch’s method per ten-second windows and the panels show means and standard error of the means for individual averages per category. Spectral powers of bilateral STNs are averaged per window. The lower row shows squared coherences between the two STNs (lower left) and between the cortex and the ipsilateral STN (lower right) and demonstrates a decrease in beta coherence and increase in cortico-STN gamma coherence during LID, with less prominent changes. Squared coherence spectra are calculated in ten-second epochs using Welch’s method. All lines were smoothed with a 3 Hz bin for visualization. Hz: Hertz, LID: levodopa-induced dyskinesia, OFF: medication Off, STN: subthalamic nucleus, sq.: squared.

### Multi-frequency cortico-subthalamic oscillatory patterns reflect dopamine-induced dyskinetic macrostate changes

Time-frequency analyses of the subthalamic and cortical recordings demonstrated changes in spectral patterns with levodopa-intake in different frequency ranges of interest (see the exemplary case in Figure 1). Spectral activity was calculated using an unbiased data-dimensionality reduction method (spatio-spectral decomposition) on the multi-channel recordings and resulted in single time-frequency data per recording site ^47^. These data are optimized independently regarding signal-to-noise ratios per frequency band of interest (theta, beta, and gamma band; see Methods). We hypothesized that distinct neural oscillatory patterns occur during hyperdopaminergic ON periods and correlate with the severity of dyskinetic periods. To this end, we first descriptively compared local STN and cortical changes occurring with the transition from OFF to ON dopaminergic periods and during different LID severity grades. All neural data from the rest and tap task were included, without specifying movement presence and absence. CDRS raters were blinded for neurophysiological results and scores were categorized into mild (less than 4 points), moderate (4 to 8 points), and severe LID (8 or more points) ^46^. Power spectral densities calculated over 10-second epochs showed an increase in subthalamic theta activity that occurred during the dopaminergic ON period without LID and gradually increased with LID-severity, whereas subthalamic low-beta activity showed a gradual decrease. Similarly, this low-beta decrease extended the low-beta decrease observed in both all non-dyskinetic periods as well as in ON-periods periods without LID after 30 minutes post-levodopa-intake (Figure 2, upper row). A large increase in narrow band gamma activity was observed in both subthalamic and cortical data predominantly during the dyskinetic periods but did not show a strong scaling effect with LID severity. Additionally, we calculated squared coherences in the same epochs to investigate how LID affects spectral connectivity between the two STNs and between the cortex and the ipsilateral STN. Inter-subthalamic coherence showed small and variable increases in theta-coherence and decreases in beta-coherence during the dyskinetic period but no obvious scaling effect (Figure 2, lower left). Gamma-activity was coherent between the cortex and the ipsilateral STN only in the two patients with severe LID.

Together, these spectral changes represent distinct neural patterns that represent medication induced macrostate changes: importantly, we describe the differences not only for the dopaminergic transition to the ON medication state without LID, but the dyskinetic state demonstrates a further spectral modulation.

### Subthalamic theta- and beta-oscillations are associated with movement absence in between dyskinetic movements

The observed changes in oscillatory cortical and subcortical neuronal pattern have been associated with LID presence in prior studies ^37,40,42^. To explore the behavioral relevance of these long-lasting, dopamine-dependent macrostate changes (Figure 2) and their interaction with short-lasting movements, we split the neural data based on the presence and absence of movement. We first defined dyskinetic versus non-dyskinetic periods based on the clinical expert assessments of video recordings of the full protocol. Dyskinetic periods were identified as behavioral states that lasted at least several minutes, and did not change on a (sub)-second resolution. Within these periods (i.e., macrostates), movement presence and absence were defined using bilateral index-finger worn accelerometer data (TMSi Saga®, TMSi International, Oldenzaal, The Netherlands). Movement absence was verified by video protocol to exclude movements on other parts of the body. Movement start- and end-points were defined on a sub-second resolution based on a thresholding procedure and led to epochs of movement presence and absence (i.e., microstates). Combining the sub second movement detection with the clinical assessments of LID presence, resulted in the categorization of all neurophysiological data into the four mentioned behavioral categories, i.e. movement presence versus absence, and dyskinetic versus non-dyskinetic periods (Figure 3). We did not differentiate between voluntary and involuntary movements during dyskinetic period, since these two concepts are not reliably distinguishable. During dyskinetic periods, patients often try to mask involuntary movements with a voluntary movement, and voluntary movements vice versa often evolve into or provoke involuntary movements. All data were merged within the respective clinically relevant behavioral categories to identify the distinct neural pattern and its modulation during microstates of movement presence. Additionally, we defined an ON condition without LID, i.e., time points after 30 minutes post-levodopa-intake without LID (n = 15), to allow comparison of dyskinetic effects versus non-dyskinetic dopaminergic medication effects. Neural data were analyzed per recording source, i.e. per STN or ECoG electrode, using the first component of an un-biased spatial spectral decomposition method to reduce the data-dimensionality while optimizing the signal-to-noise ratio ^47^. Neural data were merged within their respective microstate clusters for LID- and movement-presence and underwent time-frequency decompositions allowing for spectral comparisons per canonical frequency band of interest (Figure 4) (see Methods).

**Figure 3:**
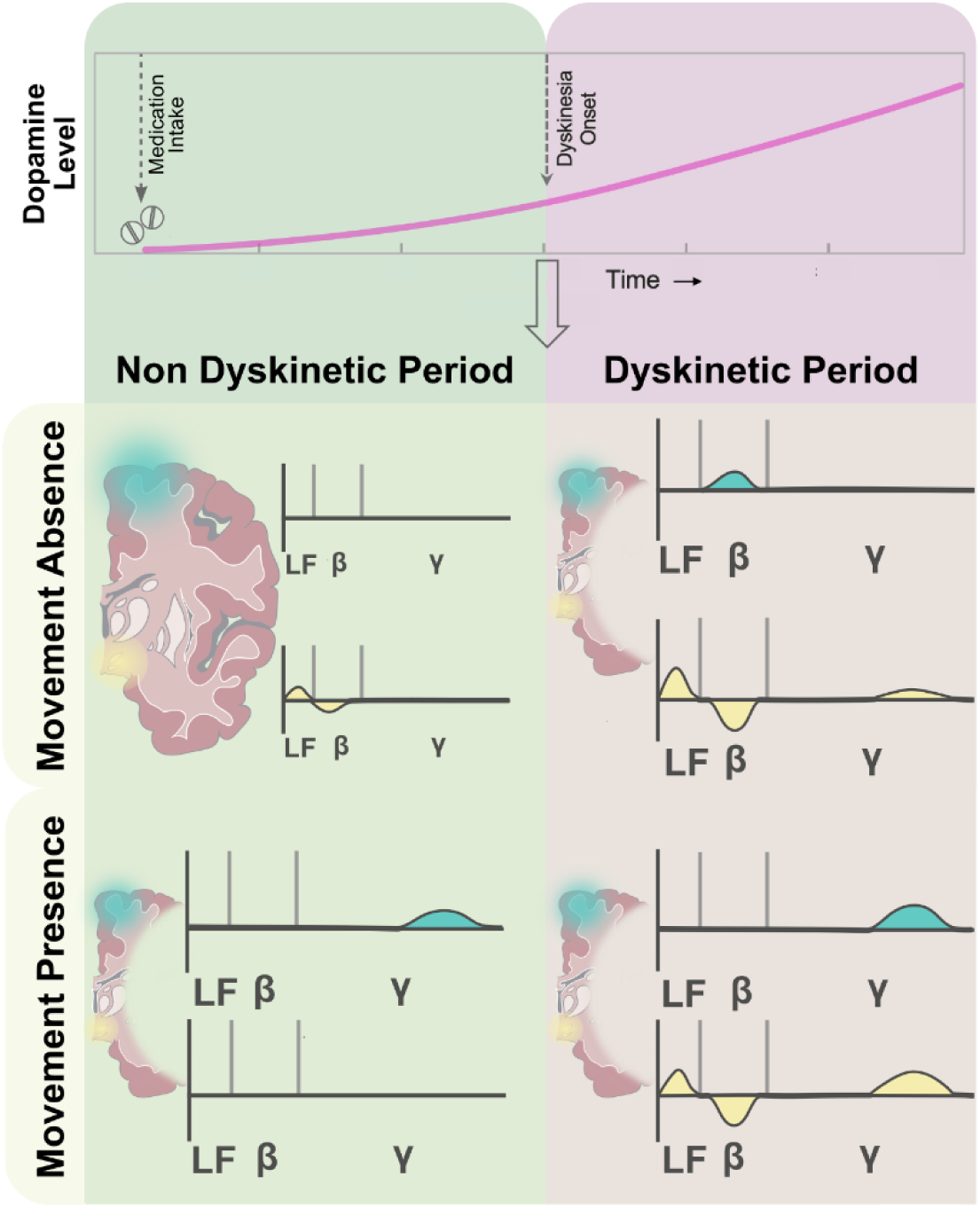
Schematic overview of dyskinesia-related spectral patterns during the presence and absence of executed dyskinetic movement. This schematic visualization illustrates that the clinical manifestation of both dyskinetic and non-dyskinetic periods contain moments of movement absence and presence. The upper line plot visualizes the rising dopamine blood-levels after medication intake. In the hyperdopaminergic state, dyskinetic periods occur when levodopa-levels are elevated sufficiently. Green PSD panels (left) represent non-dyskinetic periods, classified as a total Clinical Dyskinesia Rating Scale of zero at all time points. Purple PSD panels (right) represent dyskinetic periods. Dyskinetic periods fluctuate slowly over time with typically durations in a minute to half hour range, depending on individual disease progression, pharmacodynamics, and levodopa dosage. Movement presence and absence fluctuate Green PSDs represent cortical recordings and yellow PSDs represent subthalamic recordings. Spectral bands are shown on the x-axes: LF: low-frequency activity (theta and alpha, 4 – 12 Hz), beta activity (13 – 35 Hz), gamma activity (60 – 90 Hz). The y-axes represent the magnitude of spectral powers. LF: low-frequency, min: minutes. Spectral elevations and attenuations in this graphic are based on the spectral results, but the heights of the peaks are relative and not absolute.

**Figure 4:**
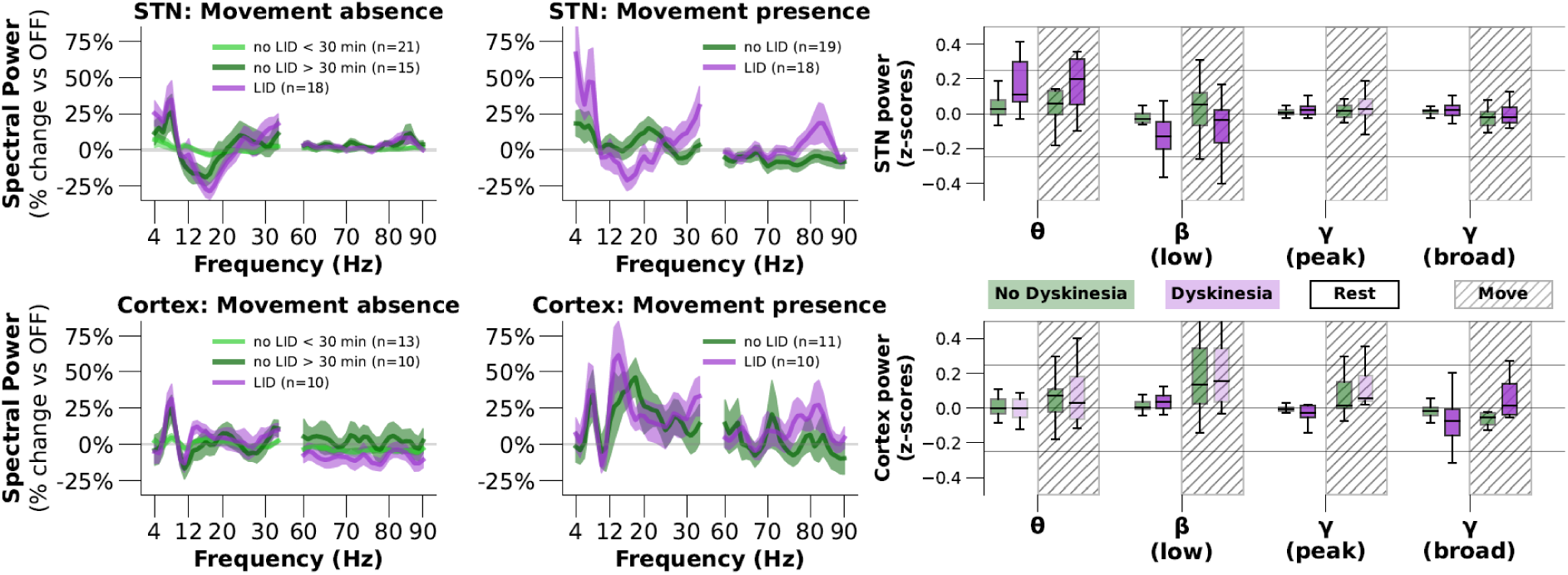
Movement modulates local subthalamic and cortical patterns during dyskinetic periods. Overall, theta increases and beta decreases in the STN were significant larger during dyskinetic states in both movement absence and presence compared to non-dyskinetic states. Increased gamma activity in the STN during dyskinetic states was mainly observed during the dyskinetic microstates with movement presence. The left four panels show power spectral densities (PSD) of subthalamic recordings (upper row) and cortical recordings (lower row). The upper and lower left PSD-panels show microstates of movement absence, upper and lower right PSD-panels show microstates of movement presence. Movement presence was assessed based on accelerometer-data according to a thresholding procedure. All green lines represent the mean over all individual average-PSDs during non-dyskinetic periods, whereas purple lines represent the dyskinetic periods. Only during movement absence, we differentiated the OFF and ON conditions within non-dyskinetic periods with sufficient sample sizes (light and dark green line respectively). Shadings show the standard error of the mean of the individual averages and legends show the number of included subjects per category. The two boxplots in the right panel visualize the mean spectral power values focused on the main spectral bands of interest (theta, low-beta, and gamma), originating from the same data as shown in the four PSD panels. For visualization purposes we decided to show these 4 main frequency bands of interest. Green and purple boxes represent respectively non-dyskinetic and dyskinetic periods. The diagonally striped backgrounds indicate movement presence data, whereas movement absence data do not have a background shading. Statistically significant differences between non-dyskinetic and dyskinetic periods, either within movement absence data or within movement presence data, are visualized as full-colored purple boxes. Transparent purple boxes indicate no significant difference compared to a neighboring green box. Alpha-values for significance testing are 0.01 and corrected for six multiple comparison (alpha is 0.0016). Hz: Hertz, LID: levodopa induced dyskinesia, min: minutes after levodopa-intake, n: number of unique subjects included in category, STN: subthalamic nucleus. Spectral band symbols: θ: theta (4 – 8 Hz), α: alpha (8 – 12 Hz, not shown), β-low: low-beta (12 – 20 Hz), β-high: high-beta (20 – 30 Hz, not shown), γ-peak: individual gamma peak (5 Hz bin around peak between 60 and 90 Hz), and γ-broad: broad-gamma (60 – 90 Hz).

Spectral power densities and squared coherences were compared between the behavioral categories, within every frequency band of interest (Figure 4 and 5). First, we describe changes in neuronal network activity that occur **within dyskinetic periods in the absence of movement execution**. *Subthalamic theta and alpha band increase and low-beta band reduction* were significantly larger in the dyskinetic period compared to the non-dyskinetic periods (resp. coefficients of linear mixed effect models (lmm-coeff.) 0.17, p=0.0001, 0.11, p<0.0001 and -0.14, p<0.0001) (Figure 4B). Importantly, these LID-related changes extended changes occurring during the ON without LID condition. When statistically testing the dyskinetic period versus the ON condition, subthalamic low-beta decrease and theta and alpha increase remained significant (resp. lmm-coeff.: - 0.07, p<0.0001; 0.09, p<0.0001; and 0.09, p<0.0001). A significant low-beta reduction also occurred in the ON versus the OFF condition, this was however a smaller effect size than the dyskinetic-related changes (lmm-coeff. -0.11, p>0.0001). Theta and alpha increases did not differ between ON without LID and OFF conditions (lmm-coeff 0.10, p=0.20, 0.08, p=0.03).

**Figure 5:**
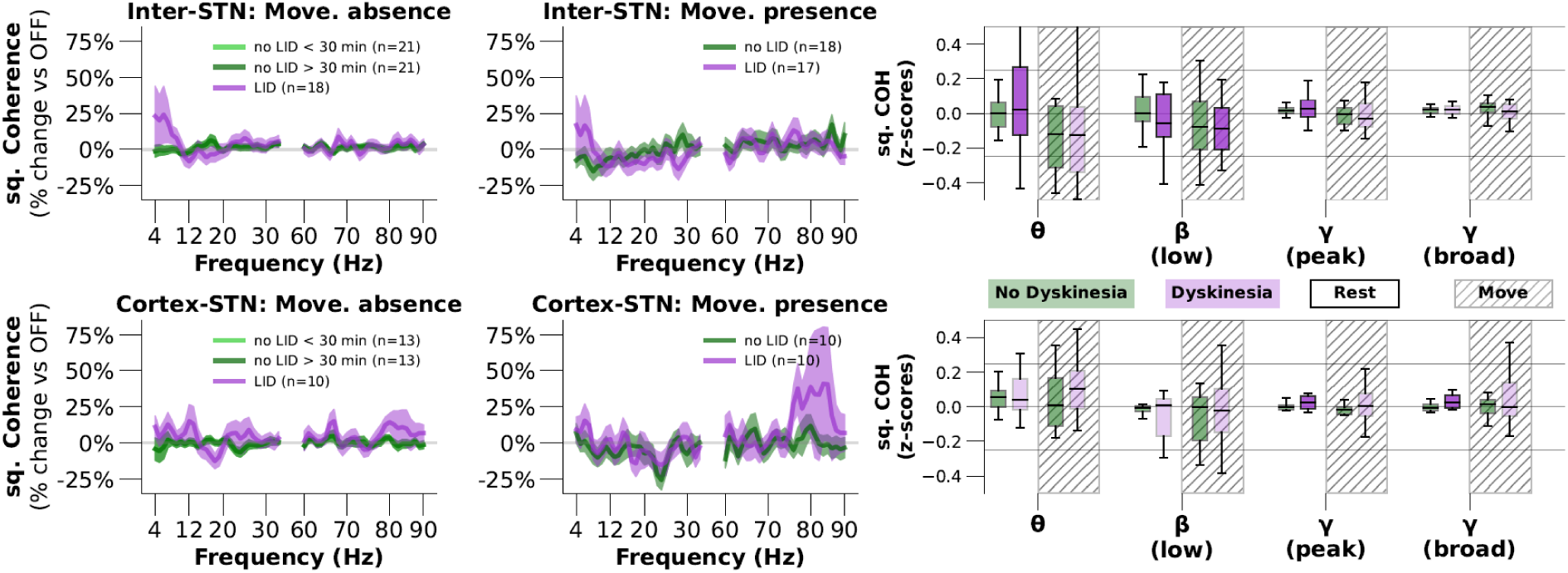
Cortico-subthalamic connectivity changes during dyskinetic periods. The left four panels show squared coherences between bilateral subthalamic recordings (upper row) and ipsilateral cortico-subthalamic recordings (lower row). The upper and lower left coherence-panels show movement absence, the upper and lower right coherence-panels show microstates of movement presence. All green lines represent the mean over all individual average-coherences during non-dyskinetic periods, whereas purple lines represent the dyskinetic periods. Only during movement absence, we could differentiate the OFF and ON conditions within non-dyskinetic periods with sufficient sample sizes (light and dark green lines, respectively). Shadings show the standard error of the mean of the individual averages and legends show the number of included subjects per category. Squared coherences are calculated based on one-second epochs using power spectral densities (psd) and cross spectral densities (csd) by Welch’s method ( │csd(source-1 and source-2)│^2^ / (psd(source-1) * psd(source-2)), see Methods). The two boxplots in the right panel visualizes the mean spectral power values focused on the main spectral bands of interest (theta, low-beta, and gamma), originating from the same data as shown in the four coherence panels. Green and purple boxes represent respectively non-dyskinetic and dyskinetic periods. The diagonally striped backgrounds indicate movement presence data, whereas movement absence data do not have a background shading. Statistically significant differences between non-dyskinetic and dyskinetic periods, either within movement absence data or within movement presence data, are visualized as full-colored purple boxes. Transparent purple boxes indicate no significant difference between a neighboring green box.Alpha-values for significance testing are 0.01 and corrected for six multiple comparison (alpha is 0.0016). Hz: Hertz, LID: levodopa induced dyskinesia, min: minutes after levodopa-intake, n: number of unique subjects included in category, STN: subthalamic nucleus. Spectral band symbols: θ: theta (4 – 8 Hz), α: alpha (8 – 12 Hz), β-low: low-beta (12 – 20 Hz), β-high: high-beta (20 – 30 Hz), γ-peak: individual gamma peak (5 Hz bin around peak between 60 and 90 Hz), and γ-broad: broad-gamma (60 – 90 Hz). Coherence plots on the left were smoothed with a 3 Hz bin for visualization.

Both individual peak gamma-activity, as well as broad-gamma activity, showed a small but significant increase during dyskinetic periods compared to all non-dyskinetic periods (lmm-coeff. resp: 0.03, p<0.0001; 0.03, p<0.0001). However, these increases did not remain significant when comparing the dyskinetic periods with the non-dyskinetic ON condition (lmm. coeffs.: 0.00, p=.98 and -0.01, p=0.04) but were significant in comparison with the OFF conditions (resp. lmm-coeff.: 0.03, p=0.001; 0.03, p<0.0001).

Regarding cortical power modulation during dyskinetic periods in movement absence, we observed small but significant low-beta increase (lmm-coeff. 0.04, p=0.0015) and gamma-activity decreases (peak- and broad-gamma resp. lmm-coeff. -0.02 and -0.08, p<0.0001 and p<0.0001) (Figure 4, lower panels). These cortical changes did not remain significant when comparing the dyskinetic periods with the non-dyskinetic ON condition (resp. lmm.-coeffs.: 0.04, p=0.03; 0.00, p=0.87; and -0.01, p=0.28). We cannot exclude that these small changes are overemphasized following the normalization influenced by the changes during movement presence that are in higher magnitudes of order.

In line with the observed STN power patterns, also *inter-subthalamic coherences significantly increased in the theta band and decreased in the low-beta band* in the absence of movement during dyskinetic periods compared to non-dyskinetic periods (Figure 5, upper row, resp. lmm-coeff. 0.35, p<0.0001 and -0.08, p<0.0001). This LID-associated theta and low-beta modulation remained significant compared to ON periods without LID (lmm.-coeffs.: 0.36, p<0.001 and -.08, p<0.001) but no difference occurred between ON and OFF conditions suggesting that the theta- and beta-coherence modulation is specific to the dyskinetic periods (lmm-coeff. resp. 0.00, p=0.935 and 0.00, p=0.855). Inter-subthalamic peak-gamma showed a minor but significant difference between dyskinetic and non-dyskinetic periods (lmm-coeff. 0.02, p=0.0004), whereas alpha and broad-gamma bands did not differ significantly (lmm-coeff. resp. -0.01, p=0.74; 0.01, p=0.08).

Cortico-subthalamic theta- and beta-coherences did not change significantly between dyskinetic and non-dyskinetic periods in the absence of movement (lmm-coeff. resp. 0.06, p=0.02; and 0.01, p=0.79), whereas alpha activity showed a trend for an increase (lmm-coeff. 0.08, p=0.001). C*ortico-subthalamic gamma coherences in peak- and broad-frequencies* showed a small but significant increases in the dyskinetic periods compared to non-dyskinetic periods (Figure 5 lower row, lmm-coeff. resp. 0.04, p<0.0001, and 0.06, p<0.0001) that remained increased compared to ON conditions (lmm-coeffs. resp. 0.04, p<0.001, and 0.06, p<0.001).

Taking together, these results demonstrate that movement absence during dyskinetic periods is characterized by increased subthalamic theta and alpha activity and decreased beta activity that significantly differ from the non-dyskinetic ON condition and thus may represent a distinct macrostate (Figures 4 and 5). In contrast, the small STN gamma power increase was more related to a general dopamine ON condition and only the increase in cortico-STN coherence was associated with the dyskinetic period.

### Dyskinetic movements are characterized by increased gamma activity in STN and cortex

To investigate whether **movement presence during dyskinetic periods** can be identified and objectified as a **microstate** in neural recordings, we isolated movements during dyskinetic periods based on accelerometer data. To ensure that network pattern changes during movement presence are specific for dyskinetic movements, we compared them with movement presence during non-dyskinetic periods, extracted from the voluntary tapping task. Subthalamic theta and alpha activity showed a trend for an increase during dyskinetic movement compared to all recorded non-dyskinetic movement (Figure 4 right upper panel, lmm-coeff. resp. 0.16, p=0.002 and 0.06, p=0.035). Low-beta activity decreased significantly during dyskinetic movement presence compared to non-dyskinetic movement presence (lmm-coeff. -0.15, p<0.001). Subthalamic broad-gamma activity, but not peak-gamma activity, increased significantly during dyskinetic movement compared to non-dyskinetic voluntary movements (resp. lmm-coeff. 0.03, p=0.0001; 0.01, p=0.044). Similarly, cortical broad-gamma activity significantly increased during dyskinetic movement compared to non-dyskinetic movement (lmm-coeff. 0.13, 0.0012, Figure 4, upper right panel). Cortical theta, beta, and peak-gamma activity did not show significant changes during dyskinetic versus non-dyskinetic movements. To examine that these changes are specific for movement during dyskinetic periods, instead of merely associated with movement during the dopaminergic ON state, we isolated all voluntary movements that occurred in non-dyskinetic states more than 30 minutes after levodopa-intake (n = 9 subjects). In line with the assumption that gamma activity codes motor parameter, peak-gamma activity showed a significant increase during dyskinetic movement compared to ON medication movement (lmm-coeff. 0.08, p=0.011), whereas theta, low-beta, and broad-gamma did not (lmm-coeffs. resp. 0.03, -0.05, and 0.0, p all > 0.12). Due to limited data availability of ON medication movement data, we only compared these main 4 frequency bands and could not assess cortical spectral changes.

Inter-subthalamic coherence during dyskinetic periods showed similar changes during movement presence as mentioned during movement absence. Theta coherence increased and low-beta coherence decreased during movement in dyskinetic periods, although only the latter reached statistical significance (lmm-coeff. resp. 0.20, p=0.006 and -0.13, p<0.001). Inter-subthalamic gamma coherence did not change significantly. Although larger cortico-subthalamic gamma coherence was observed during dyskinetic movements, none of the changes reached statistical significance, possibly due to high inter-individual variance (Figure 5, lower right).

In order to characterize the movement-related microstate changes during dyskinesia, we lastly compared the neural patterns during movement absence and presence within the dyskinetic period. Here, we found both subthalamic and cortical peak-gamma to be significantly higher during dyskinetic movement presence compared to dyskinetic movement absence (resp. lmm-coeff. 0.08, p<0.0001; and 0.24, p<0.0001). Also, cortical broad-gamma activity showed a significant increase during dyskinetic movement presence compared to dyskinetic movement absence (lmm-coeff. 0.18, p<0.001). Subthalamic broad-gamma activity however, showed a small but significant decrease in dyskinetic movement presence compared to dyskinetic movement absence (lmm-coeff. -0.02, p=0.0001). We argue that this small coefficient in the opposite direction as hypothesized, can be explained by the parallel increases in broad gamma during both dyskinetic movement absence and presence compared to non-dyskinetic periods. Modulations in cortical theta were significantly higher during dyskinetic movement presence versus absence (lmm-coeff. 0.3, p<0.001), whereas subthalamic theta and both subthalamic and cortical beta activity did not differ significantly (lmm-coeff. resp. 0.02, p=0.04; -0.01, p=0.20; and 0.06, p=0.03).

Together, when comparing movement presence with movement absence, both during dyskinetic periods, dyskinetic movement is characterized by peak-gamma increases in the STN and the cortex, and broad-gamma increases in the cortex. Subthalamic broad-gamma already shows an increase during dyskinetic periods in movement absence, which even slightly extends the increase during movement presence. These findings demonstrate that the hypothesized cortical gamma activity related to LID is specific to movement presence during LID, and that increased activity in cortical gamma as well as subthalamic peak-gamma extend gamma modulation during non-dyskinetic, dopaminergic-ON movement presence.

### Dyskinetic patterns lateralize contralateral in the cortex but spread bilateral in the STN

So far, we have observed relevant changes in spectral oscillatory activities during dyskinetic periods in both the absence and presence of movement (Table 1 and Figures 4-5). To better understand the pathophysiological role of these LID-related changes within the cortico-basal-ganglia-motor network, we investigated their laterality with respect to clinically observed LID. For this sub analysis, we selected all dyskinetic periods of unilateral LID. Unilateral LID was defined as a sum CDRS of 1 or higher for one body side, while the sum of the other body side remained 0 CDRS points. Typically, unilateral LID marks the beginning of a dyskinetic period and often occurs at the more affected side of the body. Sixteen subjects experienced unilateral LID, of which eleven received a unilateral ECoG-electrode. This led to cortical recordings from five ipsilateral and six contralateral hemispheres with respect to the dyskinetic body side and subthalamic recordings from sixteen bilateral hemispheres. Ten-second epochs during unilateral dyskinetic periods containing both movement presence and absence were selected to ensure a sufficient sample size.

STN recordings showed bilateral theta and gamma power elevation and beta power reduction during unilateral LID (Figure 6, left panel). In contrast, cortical recordings only showed narrow gamma activity in hemispheres contralateral to the dyskinetic body side (Figure 6, right panel). Cortical recordings ipsilateral to the dyskinetic body side displayed an elevated mean gamma activity without a distinct peak. Modulation of cortical theta and beta activity did not show a consistent pattern. We suggest this to be due to the limited number of included subjects per group and the lack of movement selection. The observed results suggest that the specific subthalamic oscillatory activities during dyskinetic periods occur bilateral, whereas cortical LID-related oscillatory activity occurs only contralateral to the dyskinetic body side. Based on these findings, we excluded all cortical recordings during unilateral dyskinetic periods, originating from the hemisphere ipsilateral to the unilateral dyskinetic body side for further analyses.

**Figure 6:**
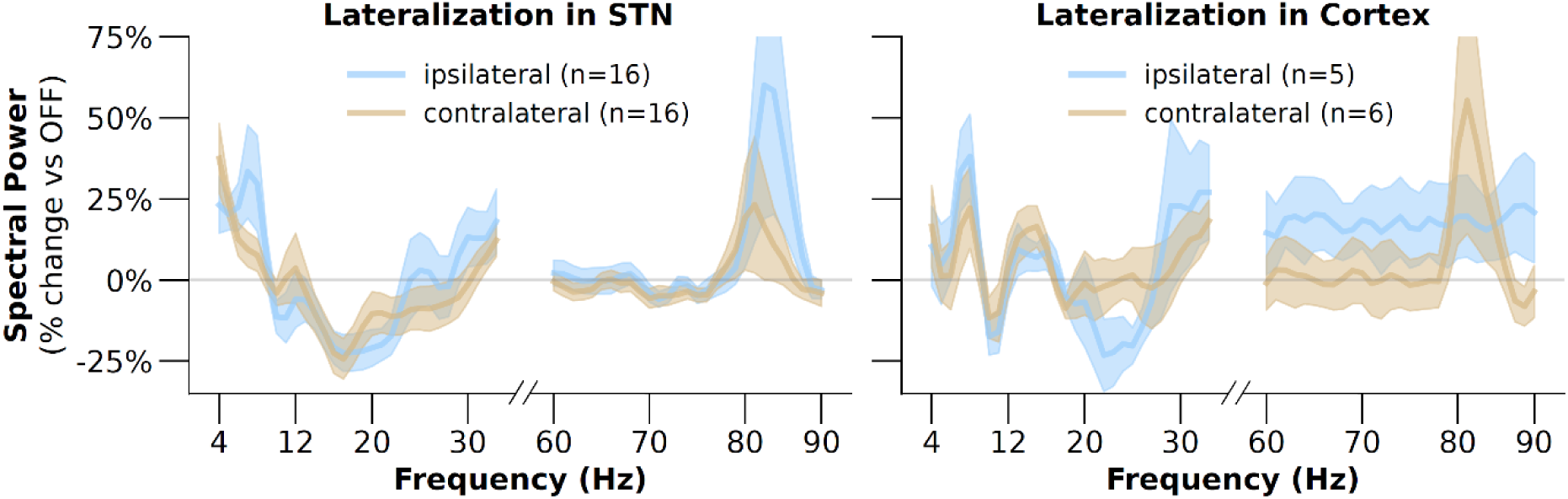
Lateralization of spectral patterns during unilateral dyskinetic periods. Percentage-changes in power spectral densities during clinical observations of unilateral dyskinetic periods are shown in subthalamic nucleus (STN) data (left panel) and cortical data (right panel). Subthalamic theta and gamma elevations and beta-reductions occur in both contra- and ipsilateral data, while cortical gamma elevations only occur in the contralateral hemisphere opposed to the dyskinetic body side. All data result from Welch-transformations per ten-second epochs, without objectified movement selection. All percentage-changes are relative to the medication “OFF” state, defined as all non-dyskinetic-moments within the first five minutes after levodopa-intake, in the absence of movement, objectified with bilateral index-finger accelerometers. The blue lines represent spectral power changes in the hemispheres ipsilateral to the dyskinetic body side; sand-color lines represent spectral power changes in the hemispheres contralateral to the dyskinetic body side. Due to the presence of bilateral STN-electrodes in all sixteen subjects with unilateral dyskinetic periods, and only unilateral ECoG-electrodes in the eleven subjects with unilateral dyskinetic periods and an ECoG-electrode, sample sizes differ per visualized group. We smoothed all lines with a 3 Hz bin for visualization. Hz: Hertz, LID: levodopa-induced dyskinesia, OFF: medication Off, STN: subthalamic nucleus.

### Subthalamic features successfully decode dyskinesia and movement-aware classification enhances predictions

To improve the neurophysiological distinction of Parkinsonian states in the context of adaptive neuromodulation, we applied our insights of distinct behavioral dyskinetic periods in several approaches predicting LID presence. First, following the consistent modulation of subthalamic theta, beta, and gamma activity during dyskinetic periods (Figures 2 and 4), we utilized each spectral band’s direction of modulation to compose one single biomarker to decode dyskinesia periods:

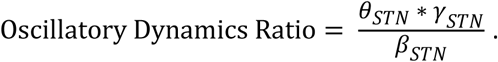

### Formula 1: Composed multi-frequency biomarker to decode dyskinesia presence based on subthalamic theta-, beta-, and gamma-power

The product of subthalamic theta power (4 – 8 Hz) and subthalamic peak-gamma power (individual gamma peak +/-2.5 Hz) is divided by subthalamic low-beta power (12 – 20 Hz). All powers are calculated over ten-second epochs using Welch’s method and are individually standardized over all recorded data. Bilateral subthalamic spectral power values were averaged per ten-second epoch and individually z-scored per frequency band over the full recording.

The Oscillatory Dynamics Ratio (ODR) biomarker (see Formula 1) leverages LID-associated subthalamic theta- and gamma-elevations and low-beta-attenuations and is expected to increase gradually during the rise of dopaminergic levels. To test the predictive potential of the ODR, all subthalamic recordings of subjects who experienced dyskinetic periods were included. All individual timelines were aligned to individual LID onset moments (Figure 7A). ODR values increased gradually after the intake of dopaminergic medication and showed a further increase after LID-onset compared to before LID onset. Then, average ODR values were calculated over ten-second windows of all 21 subjects and applied in a leave-one-subject-out cross validation. For every iteration, an optimal linear ODR threshold for LID detection was determined in each training set (n=20) and tested on the remaining test subject. Using a linear regression, the ODR could predict LID presence with a mean balanced accuracy of 0.61 (sd: 0.14), with statistically significant predictions in eight out of twenty-one subjects.

**Figure 7:**
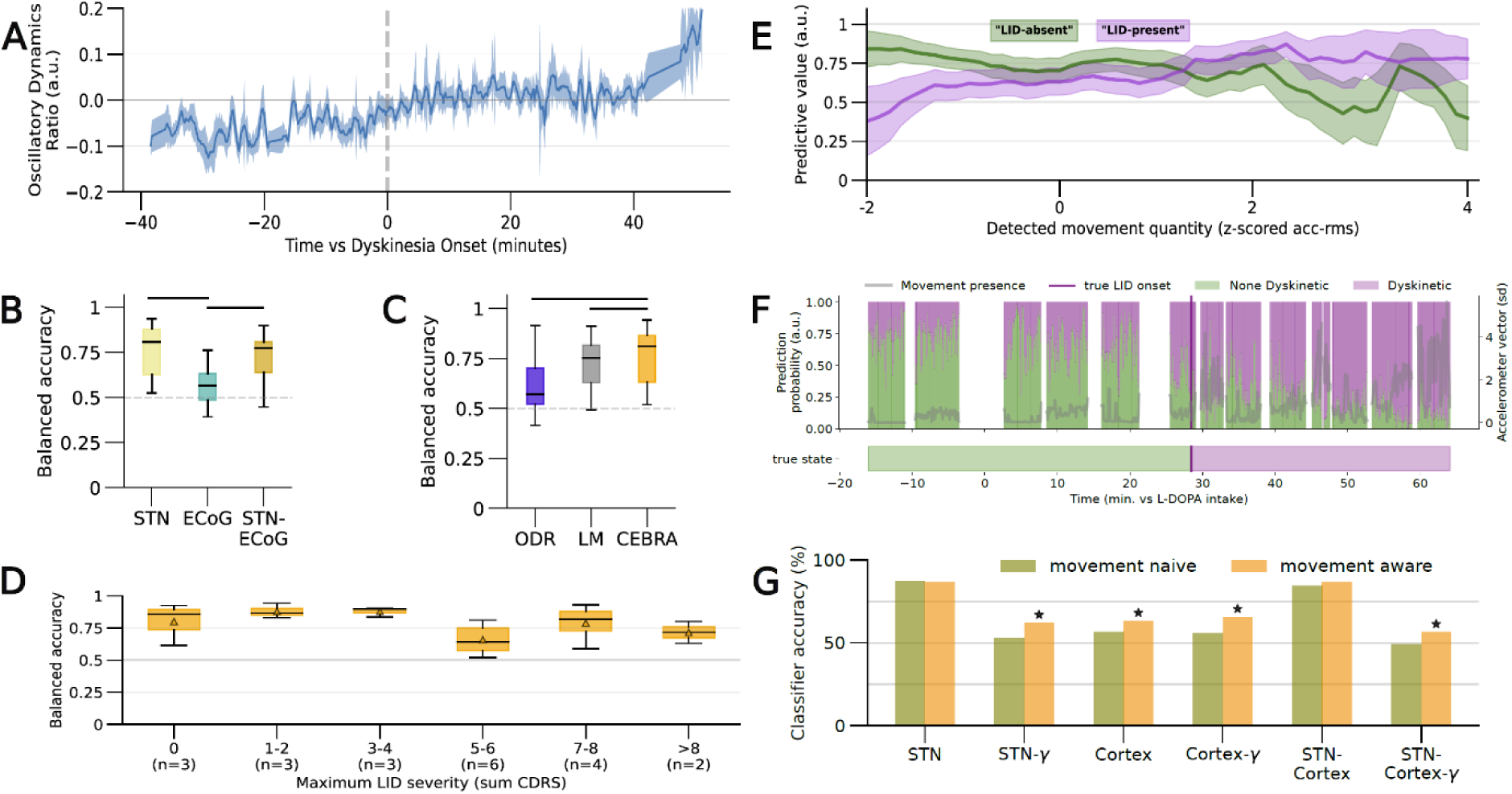
Performances of dyskinesia detection using different recording sites and models. The presence of LID was predicted using different classifier models combined with different combinations of data. Panel A shows the temporal course of a single neural biomarker, composed of the product of subthalamic theta and gamma activity divided by subthalamic beta activity (Formula 1). Ratio values were calculated per second window. The time axis is individually corrected for LID onset. Subjects without LID are not included here. The shaded area represents the standard deviation. Panel B shows the distributions of individual balanced accuracies for multivariate neural network-classifiers (CEBRA, see Methods) using only subthalamic data, only cortical data, or both. Only subjects with an ECoG electrode are included (n=13). Horizontal black lines between boxes indicate statistical difference in a paired t-test (alpha = 0.01, divided by 3 for multiple comparison correction. Panel C shows the distributions of individual balanced accuracies for classifiers using the ratio biomarker (as shown in Panel A), a multivariate linear model (STN-LM), or a multivariate neural network (STN-CEBRA). Based on superiority of the STN data, only STN data for all subjects (n=21) are included. Panel D shows the relation between the individual balanced accuracy of the LID prediction and the maximum LID severity observed within each subject. Only results of the STN CEBRA model are shown due to its superior performance. LID severity categories contain respectively 3, 3, 3, 6, 4, and 2 subjects. The horizontal black lines in the boxes represent the median and the colored triangles represent the mean of each box. The position on the x-axis categorizes the subjects on maximum LID severity as the maximum CDRS sum score assessed. Panel E shows the distribution of positive and negative predictive values (respectively PPV and NPV) over different amounts of movement objectified in the accelerometer data. Only the STN CEBRA model is included based on its highest average performance. The x-axis is categorized based on z-scored individual accelerometer root mean squares and ranges from “No movement” (z-score -2 and less) to “Many movements” (z-score +4 and higher). The lines represent the distribution of individual PPV (green) and NPV (purple) values within the respective movement quantity. The NPV values show that the reliability, or the correctness, of “no LID present” predictions, shows a decreasing trend with increasing movement presence (Spearman R = -0.07, p = 0.06). Conversely, the PPV values show that the correctness of “LID present” predictions significantly increases with increasing movement presence (Spearman R = 0.27, p < 0.001). Panel G demonstrates the prediction accuracy within the external validation subject over time. The true state bar indicates the clinically observed presence of LID and the LID onset. The upper panel shows the two prediction probabilities resulting from the STN only classifier including all three spectral bands. Probabilities for “no LID presence” (green) and “LID presence” (purple) sum up to 1.0. The grey line indicates on the right y-axis the amount of movement presence, expressed as the mean z-scored accelerometer vector magnitude. The probabilities were applied in a default binary classifier in which a probability over 0.5 lead to a positive prediction, resulting in an accuracy of 87% and an area under the receiver operator curve of 0.95. The amount of movement presence was not decisive as epochs around 10 minutes and around 50 minutes had similar movement presence, but LID was nevertheless predicted correctly. Last, Panel F shows the external validation accuracy for different feature combinations for their movement-aware versus movement-naïve classifiers. Different models were trained and tested for only STN features, only cortex features, and both STN and cortex features. For all three feature sets, one iteration was performed with only gamma-band (60 – 90 Hz) features and one with all theta (4 – 8 Hz), low-beta (12 – 20 Hz), and gamma features. All models containing subthalamic theta and beta features performed best and did not improve from a movement-aware classifier. All only-cortex models and only gamma-models were hypothesized to be more movement dependent and indeed performed significantly better with a movement-aware classifier. a.u.: arbitrary units, bal. accuracy: balanced accuracy, CDRS: Clinical Dyskinesia Rating Scale, CEBRA: neural network using latent embedding (see Methods), ECoG: electrocorticography, γ: gamma-band features only (60 – 90 Hertz), LID: Levodopa-induced dyskinesia, LM: linear model, PPV: positive predictive value, STN: subthalamic nucleus, NPV: negative predicted value.

Second, although the ODR (Formula 1 and Figure 7A) successfully differentiated between dyskinetic and non-dyskinetic periods in a sub population without requiring complex computational methods, it does not leverage the full spectrum of LID-associated neural oscillations (Table 1, Figures 4 and 5). Therefore, we tested multivariate classifiers for LID detection with different combinations of spectral features. We compared classifiers based on subthalamic features, cortical features, and cortico-subthalamic features to identify the additional value of cortical data. For each feature set, both a linear logistic regression model (LM) and a non-linear convolutional neural network using a novel latency embedding method (CEBRA ^48^) were applied (Fig S5). Since we recorded cortical data in only 13 out of 21 subjects, we first performed a head-to-head comparison of LID decoding based on only subthalamic (STN), only cortical (ECoG) and cortico-subthalamic (STN-ECoG) features in the 13 subjects with cortical recordings (Figure S5). All features were individually z-scored using the average and standard deviation from non-dyskinetic periods, in movement absence during the first 5 minutes after levodopa-intake. Both the subthalamic and cortico-subthalamic models, for both LM and CEBRA, outperformed the cortical models (Figures 7B and S5) (mean balanced accuracies: STN-LM: 0.76, sd: 0.12, STN-CEBRA: 0.77, sd: 0.14, STN-ECoG-LM: 0.75, sd: 0.12, and STN-ECoG-CEBRA: 0.72, sd: 0.15, ECOG-LM: 0.56, sd: 0.10, and ECOG-CEBRA: 0.58, sd: 0.11).

In a comparison between the ODR and the multivariate LM and CEBRA classifications, the STN-CEBRA classifier detected LID presence with a significantly higher balanced accuracy than the STN-ODR (paired t-test stat: -3.87, p = 0.001) and the STN-LM (paired t-test stat: -4.18, p = 0.001) (Figures 7C and S5).

To ensure the predictive performance of the LID classification did not depend on the severity of the dyskinetic behavior observed, we categorized all cross-validation subjects (n=21) based on their maximum LID severity, and compared the balanced accuracy distributions for different LID severities (Figure 8D). Although the small number of included subjects per category limited statistical comparisons (two to six per category), no difference in LID detection with respect to LID severity was observed.

Following our insights that the behavioral context of movement execution leads to distinct neurophysiological states during dyskinetic periods, we investigated the influence of movement presence on the LID detection in the best performing classifier (STN-CEBRA). All ten-second windows were sorted based on the amount of movement captured by the accelerometers (i.e., averaged bilateral individual z-scored accelerometer root mean squares, split in eight categories ranging from - 2 to 4 standard deviation). Based on the binary predictions within each movement presence range, we calculated individual positive and negative predictive values (respectively PPV and NPV). PPVs indicate what part of positive predictions (i.e. “LID is present”) is correct. NPVs indicate what part of negative predictions (i.e., “LID is not present”) is correct. Increased movement presence was associated with significantly higher PPVs, i.e., more reliable “LID present”-predictions (Spearman R = 0.27, p < 0.001). A decreasing trend in NPVs during epochs containing increased movement presence suggested a similar mirrored effect of more reliable “LID is not present”-detection during less movement (Spearman R = -0.07, p = 0.06) (Figure 8E).

Last, we performed an external validation on a dataset that has not been included in the main analysis (sub-024, Table S1), to demonstrate the validity and clinical translatability of our insights. We addressed the additional value of a movement-aware classifier in order to detect the dyskinetic state based on the neural oscillations described. For the movement-aware classification, we trained two separate linear classifiers on all subjects described so far (n = 21), i.e., one classifier using only 10-second-epochs exceeding a movement threshold in the bilateral accelerometer data, and one classifier using only epochs that did not exceed this movement threshold (see Methods). We tested the classification accuracy by predicting all 10-second-epochs within the external validation dataset (n = 735 epochs) with either the ‘movement-classifier’ or the ‘non-movement classifier’, depending on whether the epoch exceeded the accelerometer movement threshold or not. The resulting predictions from ‘movement-epochs’ (n = 458) and ‘no-movement-epochs’ (n = 277) were merged to calculate prediction accuracies and area under the receiver operator curve values (AUC). For the movement-naïve classification, one classifier was trained on all epochs regardless of movement presence and tested on all external validation epochs. We repeated this analysis while including only STN features, only cortical features, and both STN and cortical features. All feature combinations, i.e., mean subthalamic or cortical spectral band power, mean subthalamic or cortical spectral variation, cortico-subthalamic coherence, and inter-subthalamic coherence, were executed once including theta, low-beta, and broad-gamma features, and once including only broad-gamma features.

Statistical differences between correct predictions of two classifiers (accuracies) were compared using the McNemar’s test for binary paired outcomes. Movement-naïve classifiers including only subthalamic data from all three spectral bands reached an accuracy of 87% and an AUC of 0.95, and did not change while applying a movement-aware classification (movement-naïve accuracy STN: 87%) (Figure 7FG), similar no benefit occurred for LID detection based on STN-cortical features in all three spectral bands (movement-aware accuracy cortico-STN 87% vs. movement-naïve accuracy 84%). However, LID detection based on only cortical features, leveraging all three spectral bands, did show a significant improvement when applied in a movement-aware manner (63%, AUC 0.67 vs. 56%, AUC 0.58) (Figure 7F). Similar, all classifiers using only gamma features, i.e. subthalamic, cortical, and cortico-subthalamic features, demonstrated significantly better predictions in their movement-aware version compared to their movement-naïve version (STN: 63% vs. 53%; cortical: 65% vs. 56%; cortico-STN: 56% vs. 49%) (Figure 7F). The benefit of movement-aware classification for models based on only cortical and only gamma features underscores the close interrelation between gamma oscillation and movement on cortical level.

## Discussion

We found a specific modulation of oscillatory theta, beta, and gamma band activity in STN and over motor cortical areas that was consistently associated with dyskinesia-provoking dopamine levels, and that showed specific modulation patterns in movement presence and absence during LID in PD patients. The respective oscillatory patterns could successfully predict LID presence or absence, and movement-aware LID classification enhanced LID predictions based on both only cortical features and only gamma-band features. Here, we propose an interpretation of our findings within a framework of systemic state changes ^44^, in which we argue that the dyskinetic state as described in literature so far can be differentiated based on behavioral and neural data in two distinct state changes of which a slow macrostate depends on rising dopamine levels and a fast microstate depends on movement execution. Our insights and proposed interpretation can contribute to a better understanding of LID’s pathophysiology and can enhance currently applied clinical DBS therapies.

The systemic state principle applied follows a logic in which there is only one state at any given time for the entire system. This global state integrates many more specific sub-states, such as neural, behavioral, cardiac states and others. These sub-states can vary in their spatial and temporal dynamics, which we can acknowledge by differentiating them into microstates (short-lasting) and macrostates (long-lasting). Together, all different sub-states form an integrated system responsible for specific behavior. Using neural deep brain recordings over an entire levodopa cycle that extends the dopaminergic dosage required for clinical benefit in our patients, we were able to measure neural and behavioral sub-states within periods of levodopa-induced dyskinesia (LID).Our findings of cortico-subthalamic oscillatory patterns propose that dyskinetic periods can be interpret as hyperdopaminergic macrostates, mainly characterized by subthalamic theta elevations and low-beta attenuations, that contain distinct short-lasting microstates of movement presence and movement absence which are characterized by modulations of cortico-subthalamic gamma oscillations. These microstates fit well to the clinically observed phenotype of LID existing of the alternately execution of abnormal involuntary movements and active motor suppression.

Neural biomarkers have been identified in PD patients during dopaminergic modulation and LID ^40,49–52^, and are in line with the general pattern of oscillatory modulation of theta, beta and gamma band activity observed in our patients. Herein, increased theta and decreased beta band activity have been associated with periods of reduced motor inhibition, whereas gamma synchronization has been related to motor execution, in line with our concept of long-lasting macrostate changes and intermingled movement-specific microstate changes. Nevertheless, these temporal and behavioral aspects of neural biomarkers during LID with respect to involuntary movement periods have not been elucidated before. In order to interpret the specific behavioral relevance of the neuronal activity modulation observed during dyskinetic periods, we distinguished LID periods containing both movement presence and absence during the hyperdopaminergic state. We observed movement-independent bilateral theta increase and beta reduction of subthalamic neuronal activity that occur with the dopaminergic ON period but further increase during the dyskinetic period (Figure 4). Importantly, this slowly fluctuating macrostate change occurred bilaterally and was more prominent in periods without movements compared to movement presence suggesting that these changes are movement-independent and may rather balance motor inhibition and execution on a broader level.

Indeed, enhanced STN beta activity in PD has been characterized as a biomarker of bradykinesia and rigidity, i.e. serving as a ‘anti-kinetic’ rhythm within the cortical-STN network ^53^, that is reduced with levodopa treatment in parallel with motor improvement in PD ^54,55^. During active behavioral motor inhibition ^28^, including reactive motor stopping in the non-dyskinetic state ^43,56^, beta activity has been described to increase ^57–59^, while beta desynchronization occurs during movement preparation, - execution and -imagery ^28,55,60^ suggesting that beta desynchronization is a prerequisite change in network activity for movements to occur ^61^. In line with this concept, we observed a significant decrease in subthalamic beta activity during dyskinetic periods both during movement absence and movement presence (Figure 4), which fits previous reports of beta attenuation during LID in animals und patients ^28,40,42,49–51^. Importantly, the beta activity attenuation in dyskinetic periods significantly extended the well-known beta attenuation during the medication ON state without LID (Figure 4) ^31,55,62^ and thus, is hypothesized to represent a hyperdopaminergic macrostate change of lowered motor inhibition that facilitates involuntary dyskinetic movements. This is further supported by the significantly higher beta suppression with movement during LID as compared to movement during non-dyskinetic periods and the significant decoupling of inter-STN beta activity. The robustness of the LID-related beta attenuation regardless of movement presence, suggests that physiological and dopamine-dependent movement-related low beta synchronization and desynchronization ^43,63^ may adapt their magnitude during the hyperdopaminergic state. To further investigate the latter however, the applied data-merging method (see Methods) limits the evaluation of single movement effects. A described relation between LID severity and attenuations in cortical beta activity previously observed in rodents ^42^, was not observed in our data. This may be explained by different recording techniques, i.e. subdural ECoG recordings versus intracerebral recordings of cortical layers 5 and 6 in the rodent study.

Theta activity has been found to increase with levodopa ^64^ and during dyskinesia in PD ^37,40,50,51^. The latter has been described as a constant activity change during biphasic as well as peak-dose dyskinesia that was considered not movement-evoked ^50,51^. In line with this, we found movement-independent subthalamic theta activity increase during dyskinetic periods-underlining that the theta increase is not associated with the abnormal movement itself. Strikingly, the subthalamic theta increase in dyskinetic periods was more pronounced during movement absence. Additionally, our data show that STN theta increase correlated with LID severity (Figure 4 and S3). This is similar to observations in other hyperkinetic disorders such as dystonia, where theta activity is an established biomarker ^65^ with pallidal activity driving EMG in dystonic muscles ^66^ and increased theta power and coherence within the striato-pallidal loop correlates with severity of dystonic symptoms ^67^. On the cortical level, this relationship has so far only been described using EEG data in a few patients ^50^, but a recent ECoG study did not show improved dyskinesia prediction incorporating cortical theta activity in the model ^40^. Similar to the latter, we did not find theta modulation with dyskinesia on the cortical level, but cortico-STN coherence increased in the theta band during dyskinetic periods in our patients supporting an involvement of the cortico-STN network in this frequency band. This would also be in line with functional MRI data reporting reduction in dyskinesia during inhibitory pre-SMA repetitive transcranial stimulation, an area that was hyperactive after levodopa intake in PD prior to occurrence of dyskinesia ^45,68^.

Nevertheless, subthalamic theta elevation and increased cortico-STN theta coherence has also been described during task-related successful active movement suppression in PD ^43^ and successful suppression of involuntary tics were associated with increased cortical theta band coherence ^69^. Furthermore, elevations of subthalamic theta and inferior frontal cortex EEG power are reported during cognitive states of decision making during conflict ^37,38,70^, where STN activity has been associated with active suppression of premature responses ^71,72^. Accordingly, stimulation of the STN can increase the number of incorrect responses under conflict ^73^ or premature responses in motor inhibition tasks ^74,75^. These findings support an association of subthalamic theta activity with motor and non-motor inhibition including motor intention or involuntary movements. The functional role of theta activity seems more complex and its topographical distribution in STN recordings is more variable including non-motor ventral contacts during cognitive processing but also dyskinesia ^37,50^. Whether task-related theta modulation more prominent in the motor-cognitive domain has a different functional role as compared to theta activity recorded within the motor cortical-basal ganglia loop during hyperkinesia cannot be answered with the current data available. Our analysis suggests that an increase in theta oscillatory activity can serve as a biomarker for the slow macrostate shift towards the dyskinetic state independent of movement presence per se.

Within this macrostate of lowered motor inhibition, single dyskinetic movements occur (Figure 1) that showed an association with the increase in local subthalamic and cortical gamma activity and cortico-subthalamic gamma coherence in our patients (Figure 4 and 5). Narrowband gamma synchronization is the most established LID-related biomarker in both subthalamic ^37,40,42,49,50^ and cortical ^39–41,76^ recordings and is considered a ‘pro-kinetic’ activity ^53^. Importantly, we show for the first time a distinct movement-dependence of gamma activity associated with LID as we differentiated between movement presence and movement absence in dyskinetic periods. Previous findings on gamma synchronization during LID lack this information as current dyskinetic animal models typically lack movement absence recordings during dyskinetic periods ^77^, similar to recent naturalistic patient recordings without continuous movement monitoring on a second-scale resolution ^40^. Within our framework of macro- and microstate integrated state dynamics, we interpret these movements during dyskinetic periods as fast-changing microstates integrated in the complex phenotype of LID, which are represented by gamma elevations in bilateral STNs and the cortex contralateral to dyskinetic body sides. Movement execution has been closely associated with gamma synchronization on basal ganglia and cortical level suggesting that gamma increase may scale with the speed or vigor of voluntary movements ^35,36,60,78–81^. During a hyperdopaminergic state of reduced motor inhibition an imbalance of reduced beta activity and excessive prokinetic gamma synchronization may lead to dyskinesia. Nevertheless, it is not yet clear if narrowband gamma oscillations cause dyskinesia or are simply a physiological short-lived correlate of (excessive) movements. Moreover, recent rodent experiments suggest that excessive cortical gamma oscillations not directly trigger dyskinetic movements but propose a dysregulated cortical activity to allow involuntary movements to emerge spontaneously in downstream circuits ^82^. The latter finding may contribute to the high noise levels in our cortical data compared to our subthalamic data. In our patients, the strictly contralateral cortical gamma synchronization scaled with the dyskinesia severity and further supports that it can be used as a fast-tracking biomarker for movement activity ^40^. However, its contralateral expression opposed to dyskinetic movement makes it less robust as a biomarker in daily life and would suggest bilateral cortical recording hardware.

Subthalamic gamma activity has been argued to not merely effect movement execution, but also motor stopping ^83^. Since the clinical phenotype of LID includes active motor stopping as patients’ typically intent to terminate involuntary movements, the observed decrease in cortical gamma activity during movement absence in the dyskinetic state may also be explained in parallel to reported decreases in subthalamic and thalamic gamma activity during successful intentional motor inhibition ^43,84^.

Together, fast changing movement-dependent gamma-magnitudes could be modulated by underlying slower bilateral theta and beta rhythm fluctuations,^85^ which would comply with the here suggested framework of interpreting the complex motor network activity associated with LID into macro- and microstate changes. The movement-independence of the bilateral subthalamic theta and beta patterns supports our hypothesis that they represent a slow fluctuating macrostate of lowered motor inhibition that leads to an increased likelihood of movement to occur. The movement dependence of the STN and cortical gamma patterns suggest a more volatile and fast-changing pathophysiological role that is more lateralized within the cortico-STN network and on the cortical level. Thus, we suggest dyskinetic movements to occur as fast-changing microstates represented by gamma activity changes, parallel to a slow-changing hyperdopaminergic macrostate of lowered motor inhibition represented by a reduction in beta and increase in theta activity within the cortico-STN network.

The hypothesized macro- and microstates fit into current fundamental principles and frameworks of motor inhibition within the cortico-basal-ganglia motor network. The striatal motor network lowers its motor inhibition under chronic dopaminergic depletion by adapting the SPN-architecture ^17^ and the GABAergic ^18,20^ and glutamatergic ^19,86^ striatal projections. The hypothesized relevance of these striatal changes for the pathophysiology of LID ^87,88^, was recently confirmed by studies showing that imbalanced clusters of direct- and indirect-SPN and structural hypertrophy of SPNs allow the occurrence of LID ^22,25^. More specifically, parvalbumin-expressing (PV+) FSIs have been attributed importance in motor control ^89,90^, striatal neuronal ensembles reconfiguration ^91^, and gamma oscillation plasticity ^92^. A recently discovered PV+ selective subthalamo-striatal projection ^93^ and the presence of PV+ glutamatergic interneurons specifically in the dorsolateral STN ^94,95^, may underline the importance of PV+ FSIs in STN DBS’ sweet spot for pro-kinetic effects ^9^.

The abovementioned studies provide fundamental evidence on compensatory motor disinhibition in PD and its association with LID occurrence. Moreover, the FSI-changes may explain the characteristic gamma activity during dyskinetic movement.

Since LID is a common target for adaptive DBS ^96^, the demonstrated clinical relevance of neurophysiological biomarkers can inform future adaptive DBS (aDBS) strategies, especially to improve spatial and temporal characteristics of different closed-loop feedback mechanisms such as fast versus slow aDBS ^97^. Even more so, our external validation of movement-naïve versus movement-aware LID classification is a first demonstration of biomarkers eligible for closed-loop neuromodulation in movement disorders that uses behavioral microstates, i.e., movement presence, to enhance its neural decoding of a macrostate, i.e., dyskinetic periods (Figure 7H). The improved predictive performance with a movement-aware classifier for cortical features and only gamma features demonstrates a movement dependency for biomarkers with specific spatial and spectral characteristics, i.e., cortical patterns and gamma rhythms. Biomarkers relying on cortical oscillations and gamma rhythms therefore may be more sensible for fast, preferably movement-aware, aDBS algorithms that react to microstate changes, but may lose their specificity and effect in slow adapting algorithms. For the latter, subthalamic biomarkers may suit better using beta and theta band as biomarkers of macrostate changes. The superiority of subthalamic multivariate classifiers over the single ratio-biomarker and the cortical multivariate models (Figure 7BC, H), suggests that slow aDBS models (i.e., ten-second windows) do not profit from additional cortical features and perform best using computational heavier multivariate classifiers. The superiority of subthalamic features over cortical features to detect LID presence we hypothesize to be a result of the ten-second window length that favors slow network activity changes. We could therefore hypothesize that the prerequisite for LID to occur depends on the slower neural activity change at the level of the basal ganglia, which is in line with described discharge pattern changes within the striato-pallidal loop ^22^. Microstate changes will not be preferentially detected in our prediction model due to the selected relatively long window length that inevitably leads to the merging of microstates of movement presence with periods of movement absence within the same window for prediction. Cortical features are therefore expected to have more predictive value in fast-aDBS strategies, applying shorter window lengths, and thus isolating separate microstates of movement presence. The latter is supported by earlier work showing that cortical features had more predictive value than subthalamic features in movement prediction on a sub-second temporal scale ^35,98^.

Additionally we could demonstrate that the amount of movement presence within a ten-second window significantly affected the predictive performance of LID using an external validation data set. Our finding that “LID-present” predictions were significantly more reliable in windows containing more movement (Figure 7E) was reflected in the enhanced decoding of LID presence based on the movement-depending cortical and gamma-band features (Figure 7H). These findings underline the distinct character of the behavioral states of movement presence and absence within a dyskinetic state and suggest that future daily life LID detection models will benefit from a movement-aware classifier.

The demonstrated level of neurophysiological and behavioral detail was the result of invasive cortico-subthalamic recordings in PD patients in an in-clinic controlled environment. We addressed several analytical challenges resulting from heterogeneous individual pharmacodynamics during a LID-provoking protocol in PD patients. The before mentioned methodological challenges in rodent and naturalistic studies, i.e., a lack of movement absence during LID ^21,77^ and a lack of dyskinetic movement monitoring ^99^, underlines the importance of this translational human research to bridge the gap to valid real-world applications. Multimodal naturalistic monitoring methods such as wearables sensors and ecological valid patient reported outcomes can be applied as naturalistic proxies for kinematic and clinical observations to overcome these translational challenges ^100–102^.

Our findings propose a conceptual adaptation of neural characterizations for the dyskinetic state. Distinct spatial, temporal, and spectral neural patterns are associated with distinct clinically relevant behavioral states. A slow movement-independent macrostate change is hypothesized to occur with rising levodopa levels that is represented by bilateral modulation in theta and beta subthalamic patterns, whereas a volatile, movement-dependent microstate is associated with fast-adapting modulation in cortico-subthalamic gamma activity occurring with the execution of dyskinetic movements. The observed decoding improvement based on movement-aware classifiers for cortical and gamma features confirms these hypotheses and fits well into current pathophysiological understandings of a neural overcompensation in response to Parkinsonian motor inhibition that enables LID. Finally, this improved behavioral understanding of neural biomarkers for LID can enhance future aDBS strategies by making state classifiers behaviorally aware to optimally serve PD patients during real-life.

## Acknowledgements

We would like to thank the patients that participated in the study. We would like to thank Ulrike Uhlig for her help with organizing and assisting during the recordings. A.A.K. received funding from the Lundbeck Foundation as part of the collaborative project grant “Adaptive and precise targeting of cortex-basal ganglia circuits in Parkinsońs Disease” (Grant Nr. R336-2020-1035), from the Forschungsgemeinschaft (D.F.G., German Research Foundation) – Project-ID 424778381 – TRR 295, and from the Deutsche Forschungsgemeinschaft (DFG, German Research Foundation) under Germanýs Excellence Strategy – EXC-2049 – 390688087. P.T. received funding from the D.F.G. in Project-IDs: 433490190, 426503586, 424778381, and 426503586. WJN received funding from the European Union (ERC, ReinforceBG, project 101077060), DFG – Project-ID 424778381 – TRR 295 and the Bundesministerium für Bildung und Forschung (BMBF, project FKZ01GQ1802). J.H., R.L., and J.L.B. are fellows of BIH Charité Clinician Scientist Programs. J.H. is a postdoctoral fellow of the Parkinson Foundation.

## Author contributions

J.H., R.L., and A.A.K. conceptualized and designed the study. J.H., R.L., V.M., R.K., J.B., J.K. performed the electrophysiological recordings. K.F. and G.S. performed the neurosurgical procedures. P.K. assessed all clinical dyskinesia scores. J.H. conducted the electrophysiological, movement, and statistical analysis and drafted the study. T.B. assisted in the coherence analyses. J.H. and T.M. performed the machine learning analyses, A.A.K. and J.N. supervised these analyses. J.H., P.T., and A.A.K., contributed to the interpretation of the data, revised, and edited the work critically. J.H., V.M., and A.M. created the visualizations and performed clinical data acquisition. A.A.K. contributed critically to the supervision, project organization and funding acquisition. All coauthors have reviewed and approved the content of the manuscript.

## Methods

### Patient inclusions and characteristics

We recruited 21 patients that underwent implantation of bilateral STN DBS electrodes as a treatment for idiopathic Parkinson’s disease at the Movement Disorders and Neuromodulation Unit of the Charité University Medicine Hospital. We excluded patients with increased perioperative bleeding risks, intracranial structural abnormalities, a poor understanding of German or English language, and patients suffering from postoperative complications. This study was in agreement with the Declaration of Helsinki and was approved by the Medical Ethical Committee of Charité University Medicine (record number EA2 127 19). We described patient characteristics of our included population in Supplementary Table 1.

### Surgical protocol and invasive neural recordings

Standard clinical care subthalamic electrode implantations took place according to stereotactic gold standard procedures. All patients received Medtronic SenSight electrodes (Medtronic ©, Minneapolis, MI, US), except one patient who received Boston Scientific Cartesia X (Boston Scientific ©, Marlborough, MA, US) electrodes. 13 out of 21 patients agreed to be implanted with an additional unilateral electrocorticography (ECoG) electrode containing 6 linearly arranged contacts (Ad-tech Medical Instrument ©, Oak Creek, WI, US). To minimalize additional perioperative risks, the ECoG-electrode implantation took place at the intraoperative second hemisphere. To minimize cardiac artefacts during future chronic sensing with implanted internal pulse generators ^103^, the majority of ECoG-electrodes was implanted over the right hemisphere. All recordings took place in between the first surgery for DBS electrodes implantation and the second surgery for internal pulse generator (IPG) implantation. Details on implanted neural recording hardware is summarized in Supplementary Table 2.

### Cortical, subthalamic, and kinematic recordings

Neural and kinematic recordings were sampled via one amplifier (TMSi SAGA, TMSi International, Oldenzaal, The Netherlands) with a sampling frequency of 4000 Hz. All subthalamic and cortical local field potential (LFP) recordings had a bipolar configuration, using the lowest subthalamic ring-contact contralateral to the ECoG-electrode as a common reference. In case of no ECoG-electrode implanted, the lowest subthalamic ring-contact ipsilateral to the most affected body side was used as a common reference. A patch-electrode on the right shoulder was used as a ground-electrode. We recorded kinematic data using two tri-axial accelerometers, one attached to the distal end of each index finger. We videotaped all recordings continuously to for blinded post hoc clinical assessments.

### Behavioral protocol and clinical assessments

Patients started the experimental protocol in the “OFF”-condition, deprived from dopamine for at least 12 hours. At virtual time point zero, they took 1,5 times their dopaminergic morning dosage (100 to 300 milligrams) as fast-acting L-Dopa (Madopar LT, Roche ©, Basel, CH) (Table S1). The participants showed the first clinical signs of LID after 21.0 minutes on average (sd: 15.5 minutes). Five out of eighteen subjects that developed LID showed some first, i.e., transient, dyskinetic signs within 5 minutes after levodopa-intake. The remaining thirteen subjects had an average LID onset of 29.0 minutes (sd: 10.0 minutes). The three participants that did not develop LID received higher preoperative levodopa equivalent daily dosages (LEDD) (1873 milligrams (mg), sd: 534 mg, versus 1084 mg, sd: 455 mg, p = 0.039, Mann-Whitney-U (mwu)), and received higher experimental dopaminergic doses (250 mg, sd: 40 mg, versus 179 mg, sd: 39 mg, p = 0.037). This comparison is limited by unbalanced group sizes. The eighteen patients that did develop LID had mean peak dyskinesia severities of 5 CDRS points (sd: 3, range: 2 to 13 CDRS points), out of a maximum of 28 points (the latter corresponds to severe dyskinetic movement in all seven rated body parts).

The patients started with a 3 to 5 minutes rest recording at the beginning of the experiment around levodopa intake, which was repeated 2 to 4 times throughout the protocol. In between the seated rest recordings, the patients performed a self-paced hand tap task. Data collection continued for 60 – 75 minutes after levodopa intake and unscripted pauses were incorporated to avoid experiment abortion due to exhaustion of the patient. Data collection was continued during the unscripted pauses. During the rest recordings, we instructed the patients to sit still with opened-eyes and not talk. During the self-paced hand-tap tasks, patients remained seated and got the instruction to raise one hand and tap it one their upper leg. We instructed the patients to self-pace their taps roughly every ten seconds, without internally counting seconds. When patients paused for shorter than 7 seconds, or longer than 12 seconds in-between taps, we corrected their tapping pace.

We performed a UPDRS III assessment before levodopa intake and at the end of the recording. We collected preoperative UPDRS III scores in the dopaminergic OFF- and ON-conditions from preoperative DBS evaluations. The presence and severity of dyskinesia was assessed using the Clinical Dyskinesia Rating Scale (CDRS) throughout the whole experiment based on the video recordings ^46^. CDRS scores were assessed every five minutes, to account for fluctuations in LID severity. We chose the CDRS as clinical dyskinesia assessment based on its differentiation between LID in the upper and low extremities, and between the left and right body side. An experienced movement disorders specialist (PK) independently rated all CDRS scores, while being blinded for clinical information and neural data. To categorize LID severities in mild, moderate, and severe, we applied thresholds at 4 and 8 CDRS points. These thresholds were decided upon based on the clinical expert’s opinion after being familiar with the CDRS distribution and its correlation with clinical severity of LID.

### Kinematic signal processing and movement selection

Accelerometer time series were band passed filtered between 1 and 48 Hz (Finite Impulse Response (FIR) filter with a hamming-window) and notch filtered (FIR, notch width 4 Hz and transition width 10 Hz) to remove line noise. We down sampled accelerometer time series to 512 Hz. The filtered accelerometer signals were transformed into signal vectors magnitudes (svm) according to standard accelerometer practices ^104^. A custom movement detection was applied which combined the presence of activity and identified acceleration-peaks to categorize the accelerometer data into movement presence and movement absence. The incorporation of acceleration peaks within this categorization, helped to identify ongoing movements. Based on the variation within the svms, thresholds for movement detection were defined at 0.1 mG (milli-gravity, G: 9.8 m/s^2^) and 0.5 mG for respectively ongoing movement and acceleration peaks. The performance of this method was controlled with individual visualizations of detected movements over the course of the experiments (Figure S1 panel C and D). After confirming that the movement selection was successful for all individual subjects, the resulting time stamps of single movement start- and end-points were extracted. These time points were applied to 1) calculate the amount of movement presence within each ten-second window of neural data (as applied in the analysis showed in Figure 7), and 2) to isolate moments of movement presence and movement absence from the ten-second windows. Neural data belonging to movement presence and movement absence moments were merged separately, allowing for the neurophysiological comparison between movement presence and movement absence, as used for the analyses showed in Figures 4 and 5.

### Neural signal processing

We band pass filtered all raw multi-channel subthalamic and cortical neural time series, recorded respectively from the STN DBS electrodes and the ECoG electrodes, between 2 and 198 Hz using a FIR bandpass-filter with a hamming-window. A notch filter (FIR, notch width 4 Hz and transition width 10 Hz) removed line noise of 50 Hz including its harmonics. A threshold at eight standard deviations automatically detected segments as artefacts and visual inspections of all recordings confirmed the absence of artefacts after signal cleaning. Down sampling ensured a common sampling frequency of 2048 Hz in all neural time series.

All data were split in ten-second windows with a 50% overlap. We discarded windows with more than 33% of missing data. To reduce the data-dimensionality of the multi-channel neural recordings in an unbiased, data-driven manner, and to increase spectral signal to noise ratios, we applied the spatial-spectral-decomposition (SSD) method. One SSD transformation was calculated per frequency band of interest, on every ten-second window, including all neural channels from one neural recording source (either one STN-, or ECoG-electrode) using the Modular EEg Toolkit (https://github.com/neurophysics/meet) ^47^. SSD transforms multi-channel time series into a number of independent, singular component time series optimized for variation within a specified frequency band and uses a singular value decomposition. We selected the first SSD-component per frequency band, which holds the maximum variance within the frequency band of interest. The defined frequency bands of interest are theta (4 – 8 Hz), alpha (8 – 12 Hz), low-beta (12 – 20 Hz), high-beta (20 – 35 Hz), and gamma (60 –90 Hz). We limited the gamma range to 60 and 90 Hz based on previous literature on dyskinesia-related neurophysiological findings ^39,41,76^. We divided the selected gamma-band in three SSDs (resp. 60 – 70, 70 – 80, and 80 – 90 Hz), to prevent too large loss of broad-band activity based on the optimization based on an individual narrow gamma peak. We averaged the three broadband gamma ranges to calculate the broadband-gamma oscillations reported. The SSD-formula internally normalizes the resulting time series to the power in the neighboring frequencies (defined as ‘flanks’). To generalize this normalization step and to improve comparability of SSD results over different frequency ranges of interest, we defined the flank-frequencies for every SSD formula up to 2 Hz and 198 Hz. The only exception are the three gamma-sub-bands that were calculated with identical flank ranges between 2 and 60 Hz, and 90 and 98 Hz. The windowing and SSD transformations resulted in ten-second windows with one optimized neural time series per frequency range of interest (delta, alpha, low-beta, high-beta, and gamma-I, -II, and - III).

### Descriptive spectral analysis

Power spectral densities (PSD) were calculated with a one-second window and a 1 Hz frequency resolution using a welch time-frequency decomposition (applying a Hahn-window and a 50% segment-overlap). Only the powers within the frequency range of interest were extracted for every time series resulting from the SSD transformations (e.g., low-beta (12 – 20 Hz) time series generated the spectral power per 1-Hz bin between 12 and 20 Hz). Finally, we combined the results of all SSD time series to create a full spectrum PSD from 4 to 35 Hz and from 60 to 90 Hz, as visualized in Figures 2 and 3. To allow for inter-individual comparison and group-level analysis, we baseline corrected all spectral powers as the percentage change with respect to the OFF-condition. Using a baseline correction compared to the OFF-condition, the spectral powers contained the same temporal relevance throughout the protocol, which would be lost when a relative spectral power for each time point was calculated. Baseline corrections were performed per 1-Hz bin.

To classify spectral power changes based on the presence and severity of clinically observed LID, we labeled every ten-second window to the nearest CDRS-assessment available. Since CDRS assessments were performed every 3 to 5 minutes, to maximal distance between a spectral power value and a CDRS assessments was 1.5 to 2.5 minutes. Sum CDRS scores including all bilateral and axial items were included, except for the lateralization analysis. The latter made use of unilateral sums of both body side, excluding the axial CDRS parts in the lateralized LID score.

Squared coherence was applied as a functional connectivity metric ^52^. Similar to the spectral power calculation, we applied Welch’s methods for PSD and cross-spectral density (CSD) calculation in the formula 2. Squared coherence values were calculated between the two bilateral STNs, and the ipsilateral STN and cortex, composed from the SSD-time series and baseline corrected similarly to the spectral power values.

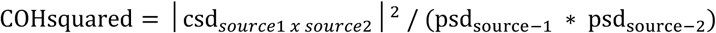

### Formula 2: Squared coherence calculation

#### Feature extraction and predictive modeling

To extract neural features for the predictive analyses, we used the SSD’d ten-second windows and calculated mean spectral power and spectral variation per frequency range of interest. Spectral powers were calculated using a Welch’s decomposition with a Hanning window and one second segments with 50% overlap. Spectral variation was calculated as the coefficient of variation of the envelope of the filtered time series (Hilbert transformation). Since SSD calculations were performed for three sub-bands of broadband gamma within 60 to 90 Hz, we included the maximum value of the three ranges as gamma value. In feature sets including subthalamic data, we included mean values of coherences between the bilateral STNs. In the cortico-subthalamic feature sets we included coherences between the ipsilateral STN and ECoG-recordings. Mean coherences values were calculated for each band width of interest using the respectively SSD’d time series, per ten-second windows.

This feature extraction resulted in individual data sets containing one value per feature for every ten-second window. Every ten-second window was labeled with the nearest dyskinesia assessment.

For the predictive analyses, we applied two different classification models, i.e., a linear model and a neural network leveraging latency embedding strategies ^48^. Both models decoded binary LID presence (0: non-dyskinetic states, 1: dyskinetic states) based on the above described features. STN models included local subthalamic features and subthalamic-subthalamic coherences. ECoG models included cortical features. STN-ECoG model included both local subthalamic and cortical features, as well as subthalamic-subthalamic and cortico-subthalamic coherences. A leave one subject out cross-validation was applied for all available subjects. For head-to-head model comparisons, only subjects with an ECoG-electrode (n=13) were included (Figure 8B). For the remaining classifiers (Panel C-E) all available subjects were included (n=21). First, we applied a linear model that decoded LID presence. For this, a ridge-regularized class-weight balanced logistic regression model, implemented in scikit-learn ^105^ (L-BFGS solver, 100 iterations) was applied. In addition to linear decoding, we utilized non-linear deep learning for classifying binary dyskinesia presence. We used the recently presented CEBRA package (Consistent Embeddings of high-dimensional recordings using auxiliary variables) ^48^. The CEBRA contrastive learning method was implemented using the “offset10-model”, a five-layer convolutional neural network implemented in PyTorch ^106^. The first layer consists of 32 hidden units, followed by three convolutional skip layers, each with 32 hidden units, and a final four-dimensional convolutional output layer. The skip layers created a bottleneck in the temporal filter dimension, reducing it from 10 to 3 samples. A Gaussian Error Linear Unit (GELU) activation function was applied to each layer, with normalization on the output layer. The model used the “constant” temperature mode, a learning rate of 0.001, and was trained for 1000 iterations with a batch size of 300.

Contrastive learning sampling was performed with the InfoNCE loss function ^107^, using a cosine distance between samples, by selecting time samples within a “time_offset” of 10, along with samples corresponding to the auxiliary dyskinesia binary training target variable. The four-dimensional embedding was following by training a ridge-regularized class-weight balanced logistic regression model, implemented in scikit-learn ^105^ (L-BFGS solver, 100 iterations), which was also applied for the non-cebra linear classifiers.

#### External validation prediction

One dataset including cortico-subthalamic recordings was used as an external validation dataset (test subject). We extracted spectral biomarkers over ten second epochs, identical to the data processing methods described above. We defined movement presence on the average of z-scored root mean square values originating from bilateral triaxial accelerometer recordings. After visual inspection, we defined a threshold of - 0.5 for movement presence classification.

For the movement-naïve classification, we trained linear discriminant analysis (LDA) classifiers on all available data from the 21 subjects reported. We applied an LDA-classifier to enable clinical translatability to future neuromodulation hardware by preventing the use of more computational demanding method such as CEBRA and convolutional neural networks. Separate classifiers were trained for the application of STN-only features, ECoG-only features, STN and ECoG-features, and for gamma-band-only features, and all spectral bands of relevance (theta, beta, broad-gamma). We tested the trained classifiers on all ten-second epochs of the external validation test subject.

For the movement-aware classification, we trained two separate classifiers for each combination of features included. We applied the same movement presence threshold for all 21 subjects as for our test subject. One classifier was trained on all no-movement epochs and tested on all no-movement epochs of the test subject (n = 277 epochs). Likewise, we trained one classifier on all movement epochs which was then applied on all movement epochs of the test subject (n = 458 epochs).

We calculated area under the curve scores on the receiver operator curve, and accuracy scores per movement-aware or movement-naïve classification. To directly compare and test statistical significance between the movement-aware and -naïve classification per combination of applied features, we applied a McNemar’s Test for two paired binary outcome-arrays ^108^.

#### Analysis software

We performed all analysis with custom-written code in Python in which we made use of standard libraries as numpy ^109^, pandas ^110^, scipy ^111^, sklearn. The custom libraries tmsi-python-interface ^112^, meet ^113^ were used for specific parts of signal processing as referred to in the methods.

## Code availability

The full code used for the processing of neural and kinematic data, performing descriptive and predictive analyses, and plotting is available on online: https://github.com/jgvhabets/dyskinesia_neurophys.

## Data availability

Data can be made available in anonymized form on reasonable request.

