## Supplemental Material for "Movement execution defines a distinct neural state in dyskinesia and enhances decoding"

**Supplementary Material** belonging to „ *Movement execution defines a distinct neural state in Dyskinesia and enhances decoding* “ by Habets et al, 2025

| Sub-ject ID | ECoG | Preop LID history | Preop UPDRS III (OFF) | Preop UPDRS III (ON) | Exp UPDRS III (OFF) | Exp UPDRS III (ON) | Max LID (total CDRS sum) | Disease duration (years) | Preop LEDD (mg) | Preop use of anti-dyskinetic and DA | Exp. L-dopa intake (mg) |
|---|---|---|---|---|---|---|---|---|---|---|---|
| **008** | **Yes** | Yes | 31 | 10 | 25 | 4 | 7 | 8 | 658 | PRAM | 150 |
| **009** | **Yes** | No | 35 | 15 | 15 | 8 | 5 | 7 | 1589 | AMAN, PRAM | 200 |
| **010** | **Yes** | Yes | 32 | 12 | 29 | 16 | 2 | 12 | 932 | AMAN, ROPI | 150 |
| **012** | **Yes** | Yes | 36 | 13 | 27 | 9 | 11 | 12 | 826 | AMAN, ROPI | 150 |
| **013** | **Yes** | No | 39 | 11 | 27 | 9 | 6 | 9 | 981 | PRAM | 200 |
| **014** | **Yes** | Yes | 64 | 27 | 32 | 16 | 2 | 14 | 1856 | none | 200 |
| **016** | **Yes** | Yes | 52 | 18 | 26 | 14 | 3 | 20 | 2276 | ROPI | 200 |
| **017** | **Yes** | No | 50 | 26 | 52 | 39 | 0 | 8 | 1415 | PRAM | 300 |
| **019** | **Yes** | Yes | 40 | 30 | 36 | 25 | 11 | 10 | 1252 | AMAN, PRAM | 200 |
| **020** | **Yes** | No | 51 | 27 | 15 | 10 | 7 | 7 | 1306 | PRAM | 100 |
| **021** | **Yes** | Yes | 52 | 30 | 39 | 20 | 2 | 10 | 905 | none | 150 |
| **022** | **Yes** | Yes | 51 | 25 | 47 | 26 | 6 | 7 | 1098 | none | 225 |
| **023** | **Yes** | Yes | 49 | 24 | 53 | 26 | 7 | 8 | 1098 | none | 250 |
| **101** | **No** | No | 51 | 23 | 41 | 32 | 0 | 6 | 2623 | PRAM | 200 |
| **102** | **No** | Yes | 41 | 11 | 13 | 16 | 4 | 6 | 513 | PRAM | 200 |
| **103** | **No** | Yes | 31 | 17 | 16 | **-** | 4 | 11 | 1408 | PRAM | 200 |
| **105** | **No** | Yes | 31 | 13 | 36 | 25 | 6 | 6 | 900 | none | 150 |
| **107** | **No** | Yes | 73 | 36 | 49 | 18 | 8 | 6 | 600 | AMAN, | 200 |
| **108** | **No** | Yes | 90 | 42 | 56 | 39 | 2 | 20 | 600 | AMAN, PIR | 100 |
| **109** | **No** | Yes | 59 | 27 | 45 | 19 | 0 | 9 | 1582 | PRAM | 250 |
| **110** | **No** | Yes | 25 | 2 | 50 | 28 | 4 | 13 | 713 | none | 200 |
| *External validation* | | | | | | | | | | | |
| **024** | **Yes** | Yes | 23 | 7 | 17 | 11 | | 10 | 1475 | AMAN | 200 |

**Table S1: Subject demographics.**

AMAN: amantadine, CDRS: Clinical Dyskinesia Rating Scale, DA: Dopamine-receptor agonists, ECoG: electrocorticography, LID: Levodopa-induced Dyskinesia, mg: milligram, preop: preoperative, exp: during postoperative experiment, OFF: OFF-state referring to dopaminergic medication, ON: ON-state referring to dopaminergic medication, PIR: piribedil, PRAM: pramipexol, ROPI: ropinirole, UPDRS III: Unified Parkinson’s Disease Rating Scale Part 3.

**Table S2: Recording details**

| Subject | STN DBS electrode type | ECoG side | ECoG electrode type |
|---|---|---|---|
| sub-008 | Boston Scientific Vercise Cartesia X | left | Ad-Tech, 1 x 6 |
| sub-009 | Medtronic SenSight | left | Ad-Tech, 1 x 6 |
| sub-010 | Medtronic SenSight | right | Ad-Tech, 1 x 6 |
| sub-012 | Medtronic SenSight | right | Ad-Tech, 1 x 6 |
| sub-013 | Medtronic SenSight | right | Ad-Tech, 1 x 12 |
| sub-014 | Medtronic SenSight | right | Ad-Tech, 1 x 6 |
| sub-016 | Medtronic SenSight | right | Ad-Tech, 1 x 6 |
| sub-017 | Medtronic SenSight | right | Ad-Tech, 1 x 6 |
| sub-019 | Medtronic SenSight | right | Ad-Tech, 1 x 6 |
| sub-020 | Medtronic SenSight | right | Ad-Tech, 1 x 6 |
| sub-021 | Medtronic SenSight | right | Ad-Tech, 1 x 6 |
| sub-022 | Medtronic SenSight | right | Ad-Tech, 1 x 6 |
| sub-023 | Medtronic SenSight | right | Ad-Tech, 1 x 6 |
| sub-024 | Medtronic SenSight | right | Ad-Tech, 1 x 6 |
| sub-101 | Medtronic SenSight | no | no |
| sub-102 | Medtronic SenSight | no | no |
| sub-103 | Medtronic SenSight | no | no |
| sub-105 | Medtronic SenSight | no | no |
| sub-107 | Medtronic SenSight | no | no |
| sub-108 | Medtronic SenSight | no | no |
| sub-109 | Medtronic SenSight | no | no |
| sub-110 | Medtronic SenSight | no | no |

**Figure S1: Graphical overview of experimental protocol.**

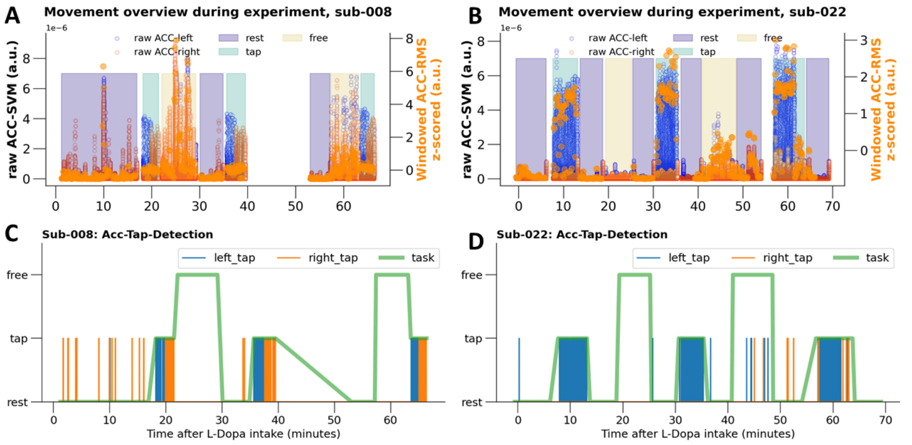

A and B show an overview of raw collected accelerometer data for left and right hands (respectively blue and orange dots) over the course of the experiment. The different background shades represent the different behavioral tasks performed. The x-axes represent the time passed since levodopa-intake in minutes.

C and D show the result of the custom movement detection method. Single movement taps are represented by the blue and orange lines for left and right hand respectively. Sub-008 (panel C) performed the tapping task bilaterally, while sub-022 (panel D) performed the tapping task only with their left hand. The little amount of taps detected during rest moments visualizes the performance of the custom tapping detection.

**Figure S2: Observed levodopa-induced dyskinesia severities**

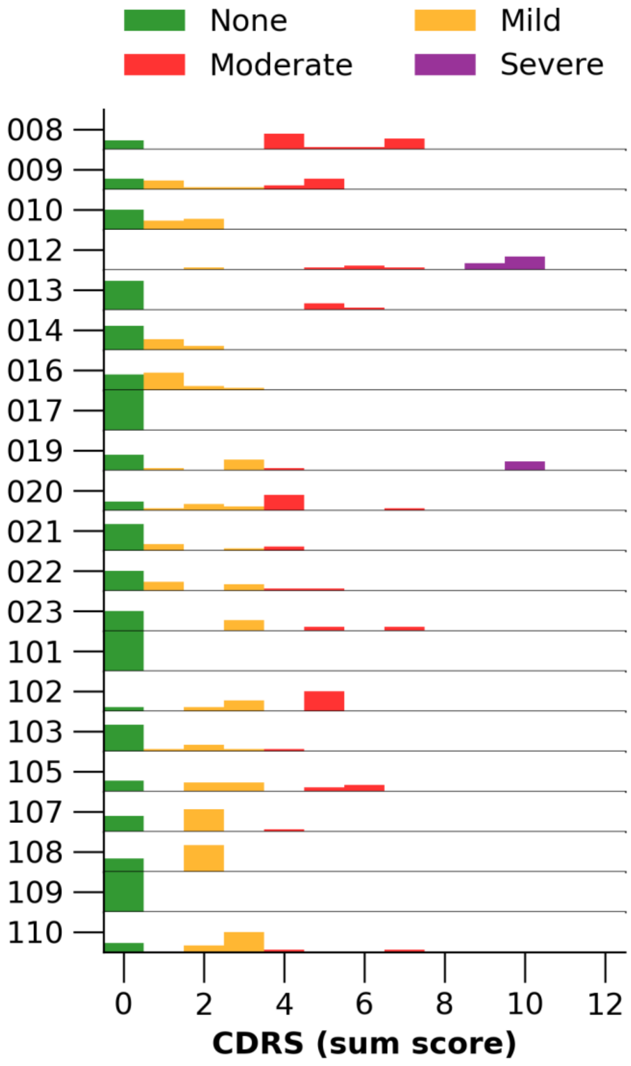

### Theta- and gamma-increase and beta-decrease scale with dyskinesia severity

We demonstrated how neuronal patterns during dyskinetic periods differ regarding the absence or presence of movement.  In this sub-analysis, we investigated the dependence of LID-related oscillatory activity on the severity of the dyskinetic periods. For this, we included the spectral powers and coherences originating from all three experimental tasks (i.e., resting states, tapping, and unscripted pauses, Figure 1) and quantified movement presence per ten-second epoch. Movement was quantified as mean root mean squares (ACC-RMS) of bilateral accelerometer data and ACC-RMS values were individually z-scored. LID severities were expressed as total sum CDRS scores.

Subthalamic theta-, beta-, and gamma-activity changed gradually with increasing LID severity (horizontal gradients in Figure 7, left column). The gradual theta increases and beta decreases occurred over the whole range of movement presence, showing no clear gradient related to the amount of movement presence (Figure 7, left, upper two panels). In contrast, gamma increases during more severe LID did show a gradient depending on movement presence. During low movement presence (i.e., ACC-RMS below 0) gamma activity only increased at high LID severities (i.e., 8 CDRS points), whereas gamma activity during higher movement presence (i.e., ACC-RMS above 0) showed increases already at lower LID severities (i.e., 4 CDRS points). These findings confirm that the LID-severity-related subthalamic gamma elevations are movement dependent whereas subthalamic theta and beta attenuations are movement independent (Figure 7, left lowest panel).

Cortical spectral patterns demonstrated similar but smaller associations with the severity of the dyskinetic period, as expected based on the cortical patterns during the dyskinetic period regardless of LID severity (Figure 2 and 4). Cortical theta activity increased and beta activity decreased gradually with increasing LID severity, without difference between the amounts of movement presence (Figure 7, right upper panel). Similar to the subthalamic findings, elevations in cortical gamma activity were only seen in mild dyskinetic periods (i.e., up to 4 CDRS points) with higher movement presence (i.e., ACC-RMS above 0). Cortical gamma in more severe dyskinetic periods (i.e., more than 4 CDS points) occurred also with less movement presence (Figure 7, right lower panel).

The observed inter-subthalamic coherences demonstrated similar patterns as the local subthalamic spectral powers (Figure S3, left column). Inter-subthalamic theta- and gamma-coherences showed a gradual increase with increasing severity of the dyskinetic period. In addition, inter-subthalamic gamma-coherence showed the strongest dependence on the amount of objectified movement presence. The coherences between the cortex and the ipsilateral STN were most clearly influenced by both the severity of dyskinetic periods, as well as the amount of movement presence in the gamma-range (Figure S3, right panels).

**Formatiert:** Englisch (Vereinigte Staaten)

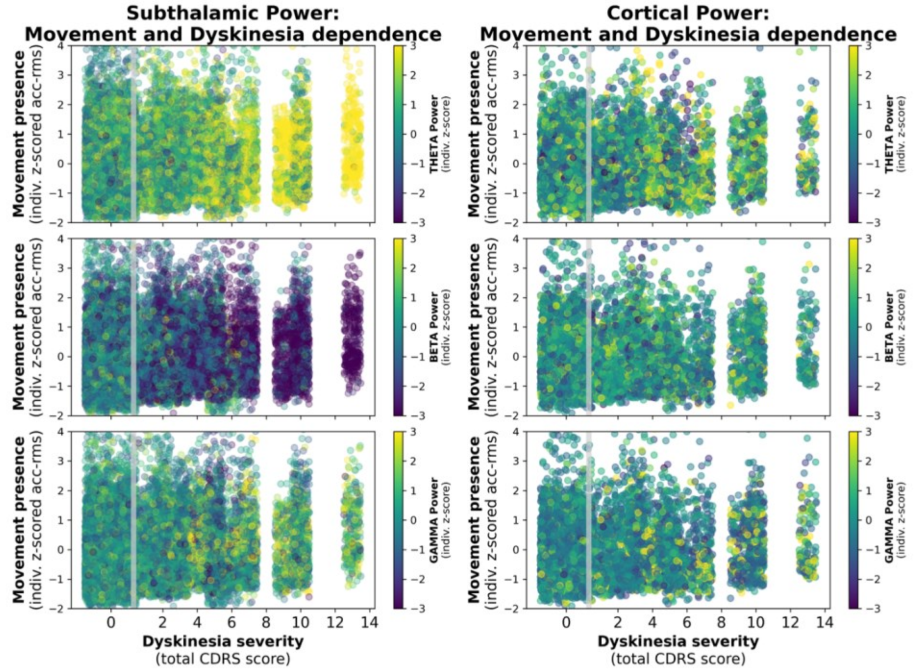

**Figure S3: Cortical and subthalamic spectral powers and their linear dependencies on dyskinesia-severity and movement presence**

The scatterplots demonstrate the modulation of theta (first row), low-beta (second row), and peak-gamma (third row) activity by dyskinesia severity (x-axis) and movement presence (y-axis). Subthalamic spectral activity is demonstrated in the left column, cortical spectral activity in the right column. Gamma activity seems to be more dependent on movement presence, whereas theta and beta modulations enhance with more severe dyskinesia. Local spectral power features were extracted from ten-second epochs of subthalamic data and averaged over bilateral STNs per epoch (left column) and cortical data (right column). Mean spectral powers are calculated per ten-second epoch (applying a 50%-overlap with consecutive ten-second epochs) and shown for the three main frequency bands of interest, i.e., theta band (4 – 8 Hz, first row), the low-beta range (12 – 20 Hz, second row), and the peak gamma range (5 Hz bin around an individual gamma-peak frequency, third row). Power values are individually z-scored and color-coded with yellow representing high values and blue representing low values. X-axes represent the absolute dyskinesia severity as total sum of the clinical dyskinesia rating scale (CDRS), including bilateral and axial dyskinetic manifestations. Y-axes represent the amount of movement present, as individually z-scored root mean squares from the bilateral accelerometer data.

**Figure S4~~3~~: Coherence features and their dependency on dyskinesia-severtiy and movement-presence**

Squared coherence features extracted from ten-second epochs of either bilateral subthalamic data (inter-subthalamic coherence, left panel) or cortical and ipsilateral subthalamic data (cortico-subthalamic coherence, right panel). Squared coherence values are calculated and data is epoched with a 50%-overlap with consecutive ten-second epochs. Mean squared coherences are plotted in the theta range (4 – 8 Hz, first row), the low-beta range (12 – 20 Hz, second row), and the peak gamma range (5 Hz bin around an individual gamma-peak frequency, third row). Coherence values

**Formatiert:** Englisch (Vereinigte Staaten)

**Formatiert:** Englisch (Vereinigte Staaten)

are individually z-scored and color-coded with yellow representing high values and blue representing low values. X-axes represent the absolute dyskinesia severity as total sum of the clinical dyskinesia rating scale (CDRS), including bilateral and axial dyskinetic manifestations. Y-axes represent the amount of movement present, as individually z-scored root mean squares from the bilateral accelerometer data).

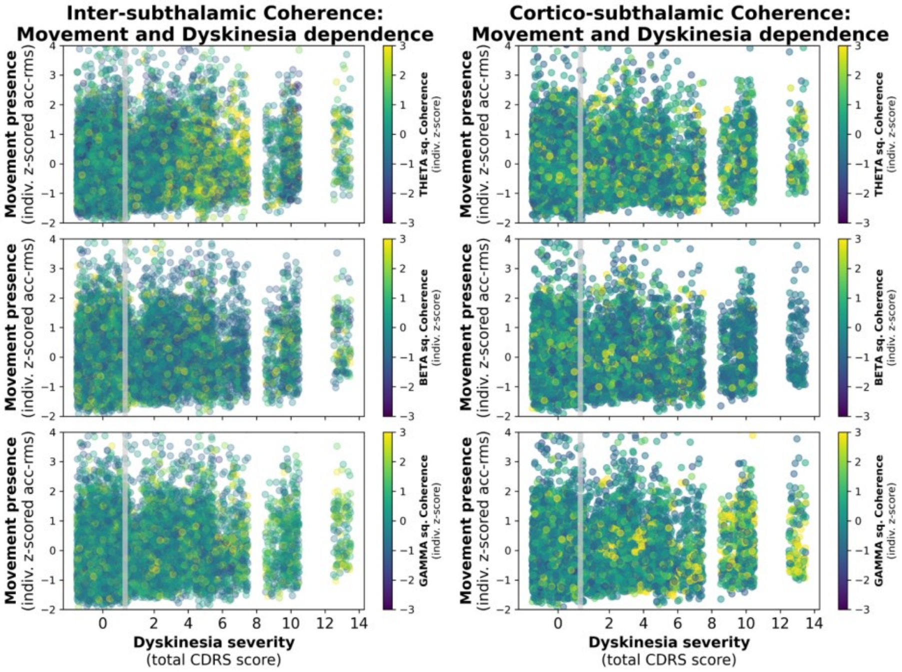

**Figure S54: Comparison of dyskinesia prediction accuracy of linear and CEBRA classifiers using STN, ECOG, or STN+ECOG features**

Both in the head-to-head comparison with identical populations (left panel), STN and STN-ECOG models were superior to ECOG models. In all feature sets, no significant difference was observed between the linear model and the CEBRA model. Including the eight STN-only subjects to the STN models did not improve the balanced accuracy (STN models right model compared to STN models left model).

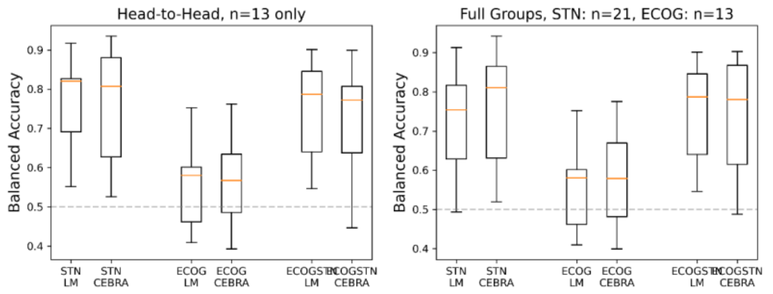

**Figure S65: Prediction performances for CEBRA models trained on two binary outcome variables for movement presence versus absence and dyskinetic states versus non-dyskinetic states.**
The left panel shows the balanced accuracy on the group level, in which the prediction model differentiated between 4 states: movement absence during non-dyskinetic states, movement presence during non-dyskinetic states, movement absence during dyskinetic states, and movement presence during dyskinetic states (random chance level prediction is therefore 0.25, indicated as the grey dotted line). Again, the STN model was superior over the ECOG model.
The right panel shows the overall accuracy for predicting the dyskinetic state correctly. Here, the movement values were disregarded in the 4-state prediction model.

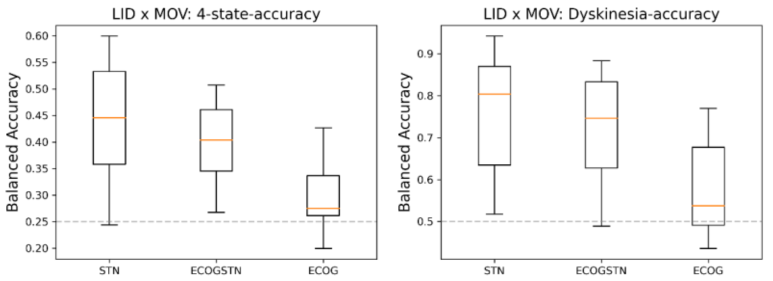
